# Systemic Inflammation Modulates Clearance and drives Extra-Hepatic Distribution of Extracellular Vesicles

**DOI:** 10.1101/2025.07.10.664102

**Authors:** Svetlana Pavlova, Doste R. Mamand, André Görgens, Antje M. Zickler, Wenyi Zheng, Xiuming Liang, Oscar P.B. Wiklander, Manuchehr Abedi-Valugerdi, Elien Van Wonterghem, Junhua Xie, Zheyu Niu, Samantha Roudi, Dongnan Yan, Guannan Zhou, Koshi Imami, Matthew P.A Wood, Roosmarjin E. Vanderbroucke, Samir El Andaloussi, Dhanu Gupta

## Abstract

Extracellular vesicles (EVs) are promising vehicles for targeted therapeutic delivery, capable of encapsulating and transporting biomolecules to specific cells and tissues. Given that inflammation is central to many acute and chronic diseases, understanding EV biodistribution under inflammatory conditions is essential for therapeutic optimization. This study examines how acute systemic inflammation influences EV biodistribution, clearance, and plasma half-life, with a focus on the role of macrophages and their polarization states. Using a lipopolysaccharide (LPS)-induced inflammation model in wild-type mice and bioluminescent and fluorescent labelling of EVs, we observed that inflammation extends the plasma half-life of EVs by over 600-fold within 2 hours and 900-fold at 24 hours post-administration, leading to significant enrichment in inflamed organs, particularly the liver and spleen. Enhanced accumulation in specific tissues translated to increased targeting of immune- and epithelial cells within those organs, with notable uptake by hepatocytes in the liver. This prolonged half-life was attributed to the altered EV protein corona in inflamed tissues, which facilitates cellular association. These findings underscore the complex dynamics between EVs and immune cells under inflammatory conditions and provide critical insights for advancing EV-based therapies in chronic inflammatory diseases.

## Introduction

Extracellular vesicles (EVs) are nano-sized membrane vesicles released by various cell types into the extracellular space and found in all body fluids^1^. They are categorized based on their biogenesis, release pathways, size, function, or composition. EVs carry and transfer biomolecules such as proteins, nucleic acids (NAs), and lipids, making them suitable candidates for therapeutic drug delivery^2^.Their capacity to cross biological barriers further enhances their potential for targeted drug delivery, making them a focus of extensive research in nanomedicine^3^.

However, the effective therapeutic application of EVs necessitates a comprehensive understanding of their *in vivo* biodistribution, clearance, and half-life. Biodistribution studies in rodent models have indicated that exogenously administered EVs are rapidly cleared from the bloodstream, with a short half-life ranging from less than 2 to 30 minutes.^4–6^ The liver and spleen act as major intravascular barriers to nanomedicine, sequestering the majority of administered EVs and preventing their distribution to other diseased tissues.^7^ Additionally, neutrophils, which constitute a significant proportion of blood leukocytes, play a pivotal role in the phagocytosis and clearance of EVs^8^.

These organs employ the mononuclear phagocyte system (MPS), a complex network of immune cells, tissue-resident macrophages, and sinusoidal endothelial cells (SECs), in coordination with various plasma proteins such as complement, to clear circulating EVs^9^. Studies have shown that the MPS system is primarily responsible for EV clearance, similar to other nanoparticle-based drug delivery vectors^10,11^. In animals with impaired innate immune and complement systems, EVs exhibit much slower plasma clearance and liver accumulation^9^. This is further supported by studies where blocking scavenger receptors on macrophages or sequestering exposed phosphatidylserine (PS) on EV surfaces prevented rapid clearance and liver uptake^12^.

Notably, a recent study identified the protein corona on EV surfaces in plasma, revealing associations with apolipoproteins and complement proteins.^13^ The EV protein corona showed significant overlap with viruses and synthetic nanoparticles, with complement opsonization enhancing phagocytosis through complement receptor-mediated uptake on macrophages.^14^ Importantly, these phagocytic cells become polarized upon exposure to inflammatory stimuli, leading to enhanced migration and modulation of their phagocytic activity.^15^ Considering inflammation as a central pathophysiology in the majority of chronic and acute diseases, understanding how inflammatory stimuli affect EV biodistribution and clearance will provide mechanistic insights into the therapeutic effects of EVs in a range of chronic disorders^16^.

This study investigates the effects of acute systemic inflammation on the uptake, clearance, and biodistribution of EVs, focusing on macrophage polarization. Using a lipopolysaccharide (LPS)-induced inflammation model in mice to simulate systemic inflammation and organ dysfunction, we track the distribution of exogenous EVs labeled with bioluminescent and fluorescent proteins conjugated to EV membrane markers. Flow cytometry is employed to analyze the molecular phenotype of cells involved in EV uptake, under both physiological and inflammatory conditions. Additionally, we examine the EV plasma protein corona to understand how inflammation affects cellular targeting. Furthermore, we evaluated the impact of enhanced extrahepatic distribution on intracellular delivery of protein therapeutics *in vivo*. By evaluating immune cell recruitment to specific organs following EV administration, we aim to provide insights into the interactions between EVs and immune cells in the context of systemic inflammation. This study sheds light on factors that influence EV biodistribution, essential for advancing EV-based therapeutic strategies in chronic inflammatory diseases.

## Materials and methods

### Cells and cell culture

HEK293 FreeStyle (FS) suspension cells expressing CD63mNG (HEK293FS:CD63mNeon Green)^17^or :CD63NanoLuc were cultured in FreeStyle™ 293 Expression Medium containing GlutaMAX (Gibco™, 12338-018) and supplemented with 1% Antibiotic-Antimycotic (A/A; Gibco™, 15240062), and 2.5 ng/mL Puromycin (Sigma-Aldrich, P8833) in 2 L polycarbonate Erlenmeyer flasks (Corning, 431280) in a shaking incubator (Infors HT Minitron) with constant shaking at 185 RPM. HEK293T-WT cells were cultured in Dulbecco’s modified Eagle’s medium (DMEM) with 10% fetal bovine serum (FBS) and 1% A/A. Bone marrow-derived macrophages (BMDMs) were cultured in Roswell Park Memorial Institute (RPMI-1640) (Invitrogen) medium supplemented with 10% FBS, 1% penicillin (100 U/mL) and streptomycin (100 μg/mL) (P/S) (Sigma-Aldrich). All cells were cultured in humidified incubators with 5% CO_2_ at 37°C. The creation and characterization of the genetically modified stable cell lines HEK293FS:CD63mNG^17,18^ were described previously. Cells were routinely checked to be free from *Mycoplasma* contamination.

### EV isolation

Conditioned media (CM) from HEK293FS:CD63mNG or HEK293FS:CD63NanoLuc cells were collected and stored at +4°C for less than 3 days until EV isolation. CM was centrifuged in two steps according to the protocol described elsehwhere^19^ to remove cells, debris, and large particles: initially, by a low-speed centrifugation step at 700 × g for 5 minutes at 4°C where the supernatant was collected for the subsequent centrifugation at 2,000 × g for 10 minutes at 4°C. The pellet was discarded, and the CM was filtered through 0.45 µm membrane vacuum filters (Corning, cellulose acetate, low protein binding, 430514) followed by filtration through 0.2 µm membrane vacuum filters (Corning, cellulose acetate, low protein binding, 430015). EVs from pre-cleared CM were further subjected to diafiltration with double volume of pre-filtered Phosphate Buffered Saline (PBS) and concentrated to the volume of about 50 mL by using tangential flow filtration (TFF) with the KrosFlo KR2i TFF System (Repligen, US) equipped with modified polyethersulfone (mPES) hollow fiber filters with 300 kDa membrane pore size (MidiKros, 370 cm^2^ surface area, SpectrumLabs, D06-E300-05-N) as a cut-off membrane at a flow rate of 100 mL/min (transmembrane pressure at 3.0 psi and shear rate at 3,700 s^−1^) as described previously.^19^ The collected samples with EVs were further filtered through 0.2 µm membrane filters. Finally, EVs were concentrated to the volume of 500 μL with the help of Amicon Ultra-15 100 kDa molecular weight cut-off (MWCO) spin filters (Amicon Ultra-15; Millipore, UFC9010 or UFC9100, respectively) at 4,000 × g, respectively. EVs were stored in 1X storage PBS-HAT storage buffer^20^ at −80°C until usage. EVs were thawed on ice prior to use^20^.

### Nanoparticle tracking analysis

The number and size distribution of particles were measured by Nanoparticle tracking analysis (NTA) ^21,22^ The analysis was performed with a NanoSight NS500 nanoparticle analyzer instrument with 3.2 analytical software and a 488 nm laser (NanoSight, Malvern Panalytic, UK). All the samples were diluted to the defined particle counts between 2 × 10^8^ and 2 × 10^9^ in 0.2-μm filtered sterile PBS. For each sample, it was recorded five 30-second videos in light scatter mode with a camera level of 14 and screen gain fixed at 2. The camera focus was adjusted until particles appeared as sharp dots. All post-acquisition settings were kept constant for all EV measurements (screen gain 10, detection threshold 5). The concentration of each sample was measured.

### Western Blot

The presence of the EV-associated tetraspanin CD63 fused to the mNeonGreen (mNG) or the Nanoluc protein, as well as the intravesical proteins Alix and TSG101, was examined by western blot. 1 × 10^10^ EVs were lysed separately using 100 µl of radioimmunoprecipitation buffer (BioRad). Then 24 µl EVs were mixed with 8 µl loading buffer (10% glycerol, 8% sodium dodecyl sulfate, 0.5 M dithiothreitol, and 0.4 M sodium carbonate). The protein was further denatured to its primary structure by incubating at 65 °C for 5 min before being loaded onto the NuPAGE+ (Invitrogen, Novex 4-12% Bis-Tris gel) and running at 120 V for 2 h. The proteins were transferred using the iBlot system (iBlot 2 Dry Blotting System; Invitrogen) for 7 min to an iBlot membrane (iBlot 2 Transfer Stacks; Invitrogen). The membrane was treated with a blocking buffer (Odyssey Blocking Buffer; LI-COR Biosciences) at room temperature for 60 min to avoid unspecific binding and later incubated overnight at 4 °C with newly prepared primary antibodies (anti-CD63 (ab134045, Abcam) diluted 1:1,000, anti-mNeonGreen antibody (EPR28835-76, Abcam), anti-Alix (MA1-83977, Thermofisher) diluted 1:1,000, anti-nLuc (N7000, Promega) diluted 1:1,000, and anti-Tsg101 (ab30871, Abcam) diluted 1:1,000). The membranes were washed 4 times with tris-buffered saline containing 0.1% Tween for 5 min each on a shaker and then incubated with secondary antibodies (goat anti-mouse (C00322) diluted 1:10,000, goat anti-rabbit (C90827-25) diluted 1:10,000) at room temperature for 1 h. The washing was repeated with one additional PBS wash, and the results were visualized using both the 700 and 800 nm channels of an infrared imaging system (LI-COR Odyssey CLx).

### Animal model

C57BL6/J WT female mice (3 months, body weight (BW) range 19 – 22 g) were purchased from Charles River Laboratories and Janvier. Mice were housed in a pathogen-free facility with free access to water and pelleted food. All of the animal procedures were performed in accordance with ethical permissions approved by the Swedish Local Board for Laboratory Animals (granted by Swedish Jordbruksverket with the number 16212-2020) and designed to minimize the suffering and pain of the animals. Blood and tissues were collected while mice were under anesthesia with 4% Isoflurane (Baxter, #VDG9623C).

### *In vivo* LPS-induced inflammation

Lipopolysaccharide (LPS) from *Escherichia Coli* 055:B05 (Sigma-Aldrich, #L6529) was dissolved to the concentration of 2.5 mg/mL in sterile PBS and stored at −20°C. LPS-induced inflammation in C57BL6/J WT mice was caused by a single dose of LPS injection at 5 mg/kg of BW intraperitoneally (I.P.) for the indicated time before EV administration. Mice were weighed before the experiment and the data were used for the determination of LPS dose for each mouse.

### *In vivo* EV administration

EVs were thawed on ice and then kept at room temperature (RT) for 10 min to warm up the solution. For *in vivo* experiments, EVs were administered at the indicated concentration by different routes: intravenously (I.V.) via tail, I.P., and subcutaneously (S.C.). Omnican-50 insulin syringes (Braun, #9151125S) were used for all EV injections. EV number for each dose was based on the NTA particle measurements.

### Perfusion of organs

For the perfusion experiment, whole blood (WB) was collected by heart puncture while mice were anesthetized with Isoflurane, and the left ventricle was infused with PBS at RT while the right atrium was perforated. 100 mL of PBS was used to perfuse one mouse. After perfusion, organs were collected in sterile PBS with 1% FBS and kept on ice till flow cytometry analysis.

### Blood collection for plasma preparation

WB from mice under terminal anesthesia with 4% Isoflurane, was collected by cardiac puncture using a 1-mL syringe into LH (Lithium heparin) tubes (BD Microtainer®, #365986) or dipotassium ethylenediaminetetraacetic acid (K2EDTA) tubes (BD Microtainer®, #365975) at RT. All the syringes and needles were pre-coated with heparin or EDTA as anticoagulant for plasma collection if the other was not specified. All the dilutions were always performed in 0.2-μm filtered sterile PBS.

### Cell-depleted plasma preparation

Cell-depleted plasma was prepared according to ISTH guidelines by double centrifugation^23,24^. There were some minor modifications due to the volume of collected blood from a mouse. Briefly, blood via cardiac puncture was collected into K2E EDTA vacutainers if another was not specified. Blood was transferred into 1.5-mL low retention Eppendorf tubes (Thermo Scientific) and centrifuged at 2,500 × g for 15 min at 4°C. The upper clear fractions were carefully transferred to new Eppendorf tubes and centrifuged for the second time by applying the same settings. This Cell free platelet poor plasma was carefully moved to new Eppendorf tubes and stored at −80°C for further measurements.

### Tissue collection for the flow cytometry analysis

After withdrawing blood by cardiac puncture, the indicated organs were immediately collected in 50-mL tubes pre-filled with ice-cold sterile PBS with 1% FBS and kept on ice till the downstream analysis. Animals were sacrificed at the determined time points indicated for each experiment. Blood was collected at the terminal time point of each experiment via cardiac puncture into K2E EDTA vacutainers at RT. In certain experiments, animals were transcardially perfused with PBS prior to organ collection to minimize contamination by plasma and blood cells.

### Tissue collection and processing for the bioluminescent assay

After the collection of blood by cardiac puncture, organs were immediately collected into 2-mL Eppendorf tubes pre-filled with ice-cold 0.1 % Triton X-100 in sterile PBS and kept on ice. The weight of collected organs was recorded. A steel ball was added to each tube and organs in 0.1% Triton X-100 in sterile PBS were stored at −80°C until analysis. Before analysis, organs were thawed on ice and homogenized at 30,000/s for 15 minutes at RT (TissueLyser, Qiagen). The samples were diluted in 0.1% Triton X-100 in sterile PBS before loading for bioluminescent assay. All data were collected in and converted to % of the injected dose (ID)/mouse.

### Bioluminescence assay on EVs, cells, plasma, and tissues using a luminometer

To quantify the level of accumulated exogenous EVs, the coding sequence for NanoLuc bioluminescence protein^25^ was fused to the sequence of the gene encoding human tetraspanin CD63, which is commonly used as a representative exosomal marker protein CD63.^26^

HEK293FS:CD63NanoLuc EV concentration was determined by the expression level of NanoLuc as Relative Luminescent Units (RLU) per EV. To quantify how many RLU contains 1 EV, 10 μl of thawed sample with EVs was diluted in 0.1 % Triton X-100 until the final volume of 25 μl at the determined concentrations in a white-walled 96-well plate. To determine the concentration of EVs in plasma and tissues, samples were diluted in 0.1 % Triton in PBS to the determined concentrations, and 25 μl was added to the same plate. The plate was shaken for 5 min at 500 RPM at RT. GloMax 96 Microplate Luminometer (Promega, US) was used to inject the substrate Nano-Glo (Promega; N1130) at 25 μl per well automatically, and luminescence intensity was recorded on a multimode plate reader.

The bioluminescence assay was used to determine the proportion of distributed EVs in the major organs, such as the liver, spleen, lungs, brain, and kidneys, and calculated as the proportion of RLU from the ID per mouse (% ID). The number of EVs in circulation was measured as % ID/mL of plasma.

### Bone marrow macrophages isolation and differentiation

Tibias and femurs were collected in ice-cold sterile PBS and bone marrow (BM) from the bone cavity was collected by flushing, using a 10-mL syringe with a G-25 needle according to the protocol.^27^ Shortly, cells were re-suspended in ice-cold PBS by pipetting and passed through 70 μM nylon mesh cells strainer to obtain a single-cell suspension. Samples were centrifuged for 5 min at 500 × g at 4°C. After removal of the supernatant, the cells were re-suspended in 1 × Red Cells Lysis buffer (ThermoFisher Scientific) according to the manufacturer’s instructions. After 2 minutes, cells were washed in PBS and re-suspended in RPMI 1640 media with GlutaMAX and Hepes, supplemented with 10 % heat-inactivated FBS, 1 % P/S. Macrophage colony-stimulating factor (M-CSF) Recombinant Human Protein (Sigma-Aldrich) at 40 ng/mL. Cells were plated in 100 mm dishes (Corning) at a concentration 2×10^6^ cells/mL in 5 mL of complete media. Cells were incubated at 37°C, 5% CO_2_ atmosphere. On day 2, 5 mL of media was added to each plate to continue stimulation. On day 4, all media was replenished with a new complete media for 3 more days. Cells were routinely checked under a microscope for cell health and confluency. On day 7, the old media was discarded, cells were washed with sterile 1 × PBS, and collected by adding 0.25% Trypsin. Collected M0 macrophages were centrifuged at 500 × g for 5 min at RT, resuspended in new complete media and counted for the experiments.

### *Ex vivo* EV uptake by differentiated mouse M0 macrophages

For EV uptake by M0 macrophages, collected M0 macrophages were plated in complete media in a 12-well plate at 1×10^6^ cells/ mL per well one day ahead to get them attached. The following day, media with unattached cells was removed, cells were washed 1 × PBS, and new media was added. Some cells were primed with LPS diluted in complete RPMI media to 100 ng/mL as a final concentration. 4 hours later 5×10^9^ EVs were added to each well for 2 hours.

### Polarization of BMDMs with LPS or IL-4 and *ex vivo* EV uptake by polarized macrophages

Differentiated mouse M0 macrophages were plated in complete media in a 24-well plate at 5e5 cells/well one day ahead to get them attached and were used as M0 differentiated macrophages for EV uptake. The following day the media was removed, cells were washed, and new media was added.

M0 macrophages were stimulated further with LPS at 100 ng/mL for 24 hours to subject them to M1 polarization or with IL4 at 20 ng/mL for 24 hours to subject them to M2 polarization. 24 hours later, 1×10^10^ EVs were added to the cells for 4 hours. Cells were always incubated at 37°C, 5% CO_2_ atmosphere.

### Flow cytometry for BMDMs

After EV uptake, BMDMs were collected by adding 0.25% Trypsin-EDTA. After 10 minutes, PBS with 1% FBS was added to each well to stop the reaction. The cells were collected by pipetting. Cellś suspension (V total 1 ml) was centrifuged in 1.5 ml Eppendorf tubes at 600 × g for 5 min and the supernatant was discarded. The cells were resuspended in PBS with 1% FBS, 2mM EDTA and transferred to a 96-well plate for staining with antibodies. Fc block reagent was added 5 min before the addition of antibodies. Cells were incubated with antibodies for 30 min at 4°C. After incubation, cells were washed with PBS with 1% FBS and centrifuged at 900 × g for 5 min at 4°C. The supernatant was discarded, cells were resuspended in PBS with 1% FBS and DAPI was added to exclude dead cells. Doublets were excluded by side scatter area versus forward scatter height gating. Cells were stained for CD11b+ (CD11b-APC/Cy7) to distinguish macrophages (F4/80-BV510), M1 polarized macrophages (CD11c-PE/Cy7), and M2-polarized macrophages (CD206-PE). EV+ cells were identified based on mNG positivity.

### Flow cytometry on cells from *in vivo* experiments

Blood from EDTA vacutainers was transferred in 1.5 mL Eppendorf tubes and centrifugated at 500 × g for 7 min at RT. The pellet was re-suspended in sterile PBS + 1% FBS by gentle pipetting at RT, transferred into 15-ml tubes, and centrifuged for the second time. The pellet was resuspended in 1 × Cell lysis buffer at RT for 10 minutes and 10 mL PBS + 1% FBS was added to tubes. The cells were centrifuged at 700 × g for 5 min at 4°C. The pellet was resuspended in PBS with 1% FBS, filtered into 5 mL Falcon tubes with cell strainer snap cap (#352235; Fisher Scientific), and stored at 4°C until the cells were stained with antibodies.

Organs (liver, spleen, and lungs) were cut into small pieces, and femur bones were crushed using a mortar for the isolation of bone marrow. Cells were separated through 70 um tissue strain and resuspended in sterile PBS with 1% FBS by pipetting. After centrifugation at 500 × g for 5 min at 4°C in 50-mL tubes, the supernatant was removed, and 1 × Red Cell Lysis buffer was added to the pellet for 10 minutes. After incubation, the tubes were filled to 30 ml with PBS with 1% FBS, and centrifuged at 500 × g for 5 min at 4°C. The washing step was repeated. The cells were resuspended in 0.5 mL of PBS with 1% FBS, filtered into 5 mL Falcon tubes with cell strained snap cap, and stored at 4°C till the cells were stained with antibodies. For flow cytometry analysis cells from the liver were diluted 1,000-fold, from spleen and bone marrow 100-fold, blood 10-fold, and no dilution for the lung cells.

The cells were resuspended in PBS with 1% FBS and transferred to a 96-well plate for staining with antibodies. Fc block reagent was added 5 min before the addition of antibodies. Cells were incubated with antibodies for 30 min at 4°C. After incubation, cells were washed with PBS with 1% FBS and centrifuged at 900 x g for 5 min at 4°C. The supernatant was discarded, cells were resuspended in PBS with 1% FBS, and DAPI was added to exclude dead cells.

The immune cell panel was changed depending on the cell type analysis. Cells from each organ were stained for a general hematopoietic cell marker (CD45-APC), macrophages (F4/80-BV510), myeloid (GR1-APC/CD11b-BV510), B cells (B220-BV510), CD4 T cells (CD45/CD3-Percp-Cy5.5), CD4-T cells (CD45+/CD3-Percp-Cy5.5+, CD4-), and neutrophils (CD45+CD11b+Ly6G+). Hepatocytes were gated as CD45-negative cells.

The data were acquired on a MACSQuant Analyzer 10 (Miltennyi Biotec). Compensation and data analysis were performed using FlowJo software. Cell populations were identified by sequential gating, and doublets were excluded by side scatter area versus forward scatter height gating. Mean fluorescence intensity (MFI) values presented refer to the expression of respective cell surface markers.

### Single-cell flow cytometry on EVs and EVs in plasma

Blood from EDTA vacutainers was transferred into 1.5 mL Eppendorf tubes and centrifuged at 500 × g for 7 min at RT. The supernatant was used for plasma isolation while the pellet was used for flow cytometry analysis on blood cells. The supernatant was centrifuged twice at 2,500 × g for 10 min at 4°C by using new tubes for every spin. Platelet-poor plasma was transferred into new 1.5 mL Eppendorf tubes and stored at −80°C until usage.

### Statistical analysis

Experimental values and counts are given as mean ± S.D. The statistical significance of differences was assessed using one-way ANOVA or two-way ANOVA with Tukey’s correction for multiple comparisons. For the comparison of two groups, paired Student’s ‘t’ test was used from GraphPad Prism 10.1.1 version. All p-values of less than 0.05 are statistically significant. Degrees of statistical significance are presented as NS > 0.05, ****P < 0.0001, ***P < 0.001, **P < 0.01, *P < 0.05.

### APEX2-Based Protein Corona Profiling

APEX2-labeled EVs were incubated with plasma from either healthy (PBS-injected) or LPS-treated C57BL/6J mice. Systemic inflammation was induced via I.P. injection of LPS (5 mg/kg) 4 hours prior to blood collection. EVs were incubated with 100 μL plasma at 37°C for 30 minutes, followed by addition of biotin-phenol (500 μM) and H₂O₂ (1 mM) for 1 minute to initiate APEX2-mediated biotinylation. The reaction was quenched with sodium ascorbate, Trolox, and sodium azide. Biotinylated proteins were enriched using streptavidin-conjugated magnetic beads and processed for on-bead digestion using trypsin. Peptides were analysed by LC-MS/MS (Orbitrap Eclipse mass spectrometer, Thermo Fisher Scientific) and data were searched against the mouse UniProt database using MaxQuant.

Raw protein intensities were filtered to remove common contaminants (cRAP) and red blood cell (RBC)-associated proteins. Protein abundances were normalized across replicates and differentially enriched proteins (log₂FC ≥ 1 or ≤ –1; adjusted p-value ≤ 0.05) were identified using the Limma package in R. Gene Ontology (GO) and Reactome pathway enrichment analyses were performed using clusterProfiler and ReactomePA.

### *In Vivo* Functional Delivery Using Ai9 Cre-Reporter Mice

Cre recombinase protein loaded EVs were generated by transfecting HEK293T cells with plasmid expressing CD63 Intein Cre, EVs were purified as described earlier. Ai9 mice (Jackson Laboratory, Stock #007909) containing a loxP-flanked STOP cassette upstream of the tdTomato reporter were used to assess *in vivo* Cre delivery. Mice were divided into four groups (n = 3–4 per group): PBS only, LPS only, PBS + CD63 Intein Cre EVs, and LPS + CD63 Intein Cre EVs. Systemic inflammation was induced as described above. EVs (1×10^12^ particles in 200 μL PBS) were administered I.V. 4 hours post-LPS injection. After 7 days, mice were perfused with PBS, and tissues (spleen, liver, kidney) were harvested, fixed in 4% PFA, and cryosection.

Fluorescence microscopy was performed to detect tdTomato expression, indicating successful Cre-mediated recombination. DAPI was used to counterstain nuclei. Signal intensity was compared across groups. No increase in tdTomato signal was observed in EV-treated groups, consistent with inefficient endosomal escape of EV-delivered Cre. Background tdTomato fluorescence was noted in spleens of PBS and LPS-only mice, possibly due to leaky reporter activity or autofluorescence under inflammatory stress.

### Single EV Imaging Flow Cytometry

Blood from EDTA vacutainers was transferred to 1.5 mL Eppendorf tubes and centrifuged at 500 × g for 7 min at RT. The supernatant was used for plasma isolation while the pellet was used for flow cytometry analysis on blood cells. The supernatant was centrifuged twice at 2 500 × g for 10 min at 4°C by using new tubes for every spin. Platelet-poor plasma was transferred into new 1.5 Eppendorf tubes and stored at −80°C till was used. For the analysis of EV at the single vesicle level, high resolution Imaging Flow Cytometry (IFCM) on a Cell stream instrument (Cytek/Amnis) based on previously optimized settings and protocols established on an Amnis Cellstream instrument (Amnis/Luminex).EVs from plasma were diluted in PBS-HAT buffer (ref 4) in all steps. In brief, 2.5 ×10^8^ EVs (NTA-based particles) were stained in a total volume of 25 µl with 8 nM of fluorescent antibodies against human CD63 or human IgG isotype control antibodies (Miltenyi Biotec). EVs were incubated overnight at room temperature in the dark with a subsequent dilution to a concentration of 1×10^7^ EVs/ml in a final volume of 100 µl before data acquisition. Unstained samples and non-EV-containing samples incubated with antibodies were included as controls, respectively.

## Results

### EVs display distinct pharmacokinetic properties *in vivo* in an inflammatory state

Inflammation is known to play a crucial role in regulating macrophage activity, influencing their polarization and ability to engulf extracellular materials, including EVs^28,29^. During an inflammatory response, macrophages undergo functional changes that can enhance or suppress their phagocytic activity, depending on their polarization state (M1 pro-inflammatory or M2 anti-inflammatory)^30^. This modulation in macrophage function is likely to impact the uptake of EVs, as macrophages are primary phagocytes that actively participate in clearing or distributing EVs throughout the body. Given this relationship, we aimed to investigate how acute inflammation might affect the biodistribution and clearance of EVs in vivo.

To investigate this, we utilized our previously optimized bioluminescence labelling strategy by stably expressing CD63-NanoLuc fusion in the EV producer cell line^31^. In order to minimize variability, the HEK293FS cell line was used exclusively as the platform for EV production.

EVs purified from HEK293 CD63-NanoLuc FreeStyle cells, exhibiting typical EV characteristics of 90-110 nm in size and presence of EV markers CD63, ALIX and TSG101 (Figure S1), were injected into animals pre-treated with LPS to induce systemic acute inflammation. EVs were administered via different routes, including I.P. and I.V., to evaluate the impact of inflammation on their biodistribution. Blood and tissue samples were collected at 30 minutes, 2 -, 4 -, and 24 hours post-administration to assess EV accumulation in different organs (Figure 1a).

**Figure 1.**
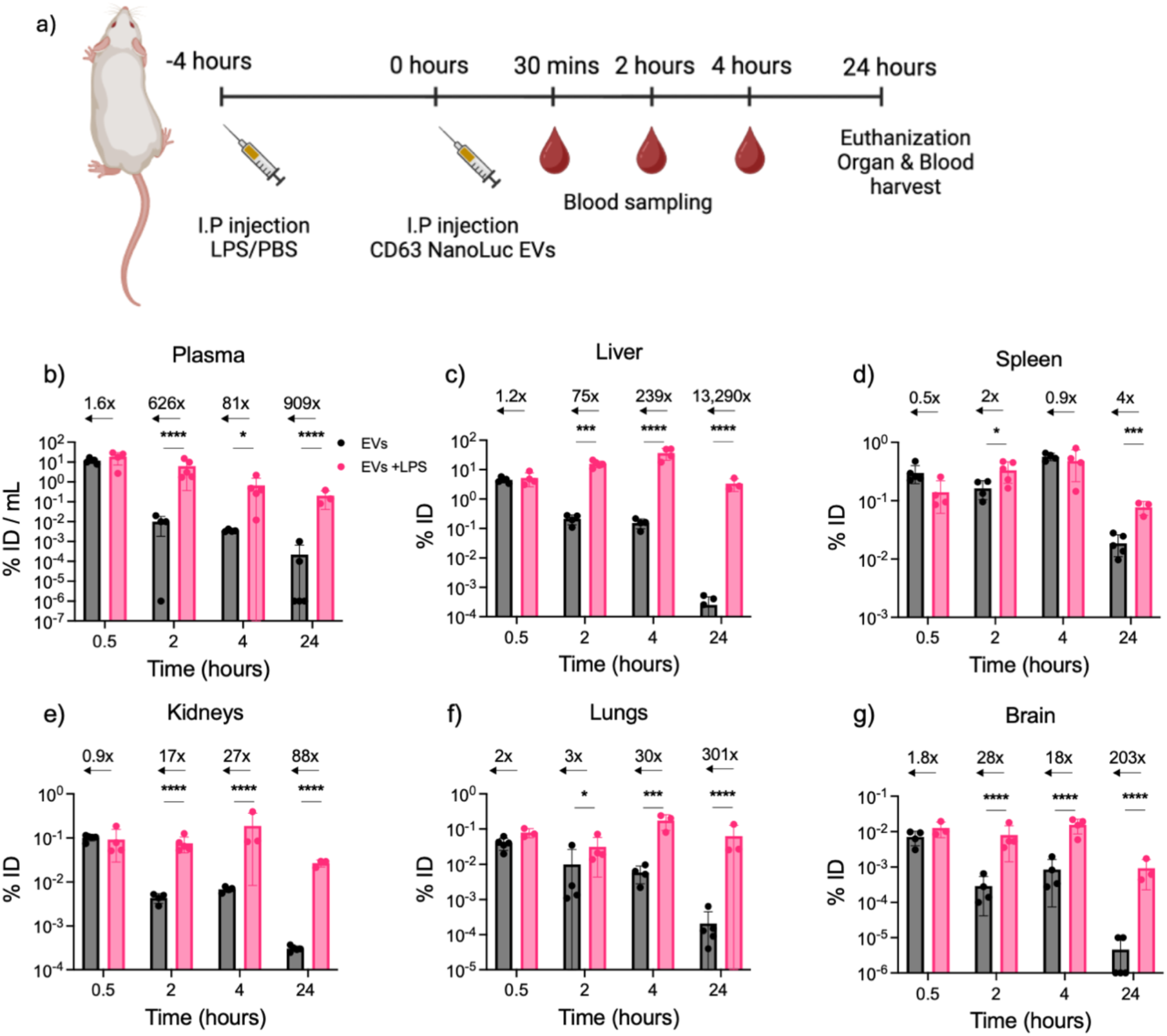
I.P. administered EV biodistribution in tissues. (**a**) Schematic workflow for studying biodistribution of exogenous EVs *in vivo*. (**b – g**) EV accumulation in the respective tissues was quantified as % of injected dose - % ID (organ/mouse) or % ID /mL of plasma employing the bioluminescent assay. The numbers display RLU fold change in EV accumulation in LPS-primed mice over the healthy mice in the brain (**b**), lungs (**c**), liver (**d**), spleen (**e**), kidney (**f**), and plasma (**g**). The acute inflammation was induced by a single dose of LPS at 5 mg/kg 4 hours before the EV administration. Based on NTA measurements, EVs were injected I.P. at 1×10^11^ EVs/dose (n = 4 – 5 mice) for 0.5, 2, 4, and 24 hours before the blood and tissue collection according to the Methods section for the subsequent analysis. A control group was used for the normalization of the data (n = 4). Black – non-primed healthy mice, pink – LPS-primed mice. Arrows indicate the fold change in EV accumulation in LPS-primed mice over healthy mice (primed with PBS). Statistical analysis was performed by two-way ANOVA. ** represents p<0.01; ***, p<0.001; and ****, p<0.0001. The results represent mean ± SD.

Our findings revealed that within 30 minutes, the proportion of circulating EVs in LPS-primed mice was 1.5 times higher compared to healthy controls (18.6% vs. 11.8% Injected Dose of the injected dose (ID), respectively) (Figure 1b). This fold increase in plasma half-life grew substantially over time, with exogenous EV levels in plasma increasing 626-fold at 2 hours post injection and over 909-fold by 24 hours post injection in LPS-inflamed mice (Figure 1b). The extended retention and distribution of EVs in inflamed mice suggest a delayed clearance process, likely influenced by changes in macrophage activity and other immune cell functions during inflammation.

EVs predominantly accumulated in the liver and spleen under both healthy and inflamed conditions (Figure 1c, d), with LPS-induced inflammation leading to much higher EV retention. In LPS-primed mice, EV accumulation in the liver increased by 10.6% ID between 30 minutes and 2 hours, while in healthy mice, liver EV levels decreased by 21-fold during the same period (Figure 1c). After 24 hours, the difference between the two groups reached 13,290-fold (Figure 1c). Notably, EV levels in the spleen showed no major differences between LPS primed and healthy mice (Figure 1d).

Moreover, at the 2-hour mark, EV elimination was significantly delayed in LPS-primed mice, showing a 17-fold increase in the kidneys (Figure 1e), a 3-fold increase in lungs (Figure 1f), and a 28-fold increase in the brain (Figure 1g), compared to healthy mice. These findings suggest that inflammation not only prevents EV phagocytosis but also delays the clearance of EVs from various tissues, leading to prolonged retention. This is particularly evident in organs rich in immune cells, such as the liver and spleen.

Next, we explored whether the timing of inflammation induction influenced EV distribution across tissues. LPS was administered either 30 minutes or 4 hours prior to EV injection, or concurrently with EVs, and tissues were harvested 4 hours after EV treatment (Figure 2a). In several organs, we observed a direct correlation between the induction timing of inflammation and EV distribution. For example, in the liver, EV accumulation increased from a 66-fold difference as compared to healthy mice (when EVs were injected simultaneously with LPS) to a 239-fold difference when EVs were administered 4 hours after inflammation induction (Figure 2b). Similar trends were observed in the brain (Figure 2c) and kidneys (Figure 2d), though the magnitude of the change was lower.

**Figure 2.**
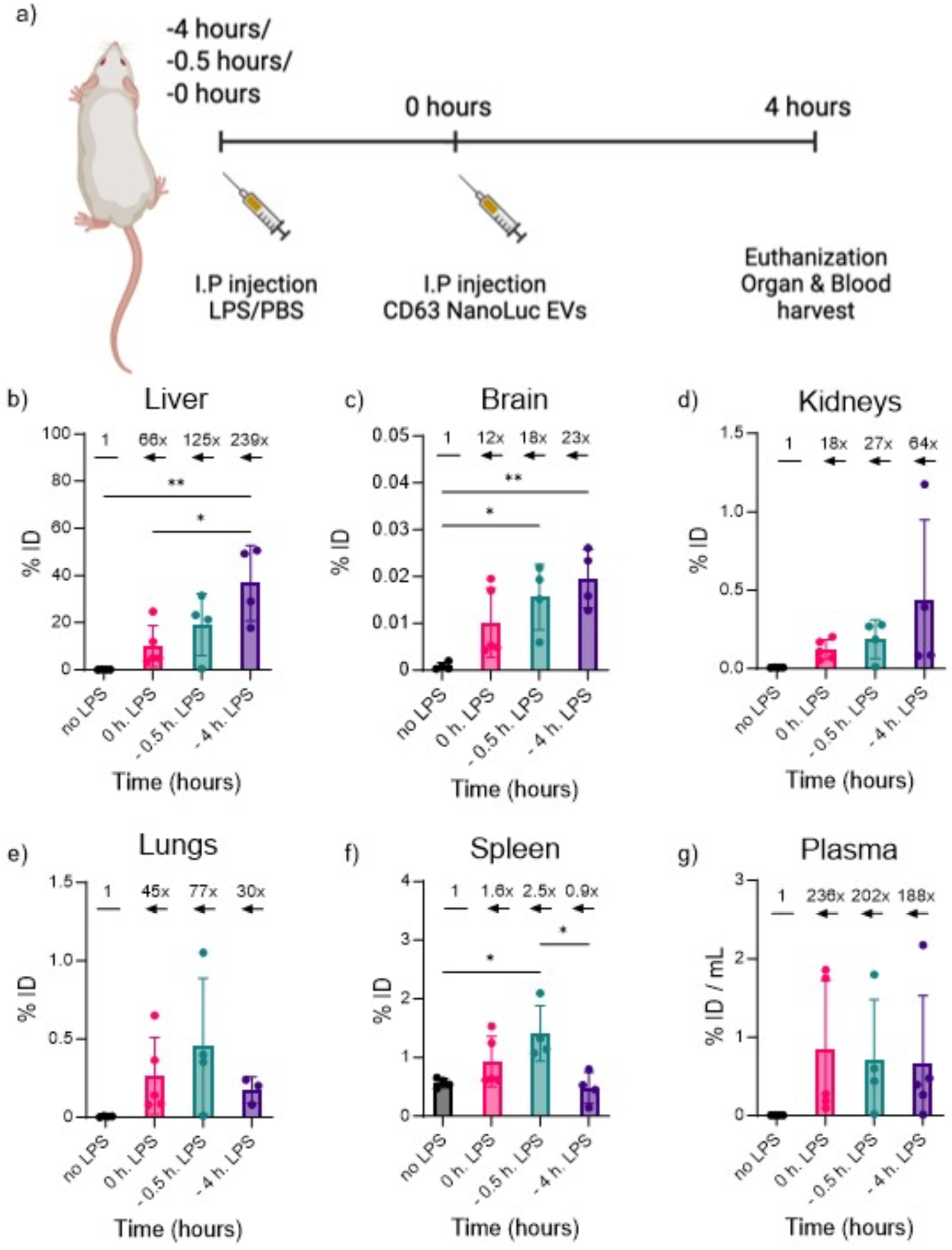
Correlation between EV accumulation in various organs and the time of LPS-induced inflammation. (**a**) Schematic overview. (**b – g**) EV distribution quantified by the bioluminescent assay as % ID (organ/mouse) in the Liver (**b**), Brain (**c**), Kidney (**d**), Lung (**e**), Spleen (**f**), or % ID /mL of plasma (**g)**. The numbers indicate RLU fold change in LPS-primed mice over the healthy mice. The acute inflammation in mice was induced by a single dose of LPS at 5 mg/kg (I.P.) at the same time with EVs (0 hours), 0.5 hours, or 4 hours before the administration of EVs. 1×10^11^ HEK293FS:CD63NanoLuc EVs/mouse were injected I.P. (n = 4 – 5 mice). WB and organs were collected 4 hours after the EV injection. A control group was used for the data normalization (n = 2). Black – only EVs; pink – LPS and EVs were injected in parallel; cyan – EVs were injected 30 min after priming with LPS; violet - EVs were injected 4 hours after priming with LPS. Statistical analysis was performed by one-way ANOVA. * Represents p<0.05, **represents p<0.01. The results represent mean ± SD for % ID or mean ± SEM for fold change graphs.

In contrast, the highest EV accumulation in the lungs and spleen occurred when inflammation was induced 30 minutes before EV administration (Figure 2e, f). In all inflamed mice, circulating EVs were still present, 4 hours post EV injection, at low but consistent levels (0.67-0.8% ID), whereas EVs were nearly undetectable in healthy mice (0.004% ID) (Figure 2g). These data underscore the importance of inflammation onset in modulating EV distribution patterns. Finally, we assessed how the route of administration and the severity of inflammation impacted EV biodistribution. LPS was administered at two doses (2.5 and 5 mg/kg) and EVs were injected S.C. Plasma EV levels remained low (<0.03% ID) with no significant differences among groups (Figure S2). However, in tissues such as the liver, lungs, and kidneys, EVs were detectable at a lower level and the levels correlated with the LPS dose. (Figure S3). Interestingly, no EVs were detected in the spleen of LPS-primed mice, whereas EVs were present in the spleen of healthy controls (Figure S3). Importantly, levels of EV enrichment in organs were lower as compared to systemic injection, similar observation is also previously reported by us and others, where S.C administration fails to show a significant systemic distribution^31^.

When EVs were administered I.V., they were no longer detectable in the plasma of healthy mice after 4 hours. However, in LPS-primed mice, EVs were still present at 0.5% ID (Figure S2). This suggests that LPS-induced inflammation significantly prolongs the presence of EVs in circulation and delays their clearance from the body. Overall, our data demonstrate that inflammation has a profound impact on EV biodistribution and clearance. The disease severity, timing, and route of EV administration all contribute to these changes.

### EV uptake is dependent on the activation status of macrophages

To assess how macrophage activation state influences extracellular vesicle (EV) uptake, bone marrow-derived macrophages (BMDMs) were first differentiated into the M0 state using M-CSF and then polarized into M1 (LPS-treated) or M2 (IL-4-treated) macrophages. CD63-mNeonGreen (mNG)-labelled EVs were added for 4 hours, and flow cytometry was used to evaluate surface marker expression and EV internalization.

High CD11b expression was consistently observed in EV+ macrophages across all polarization states (3a), indicating that CD11b high cells are the primary EV-uptaking population regardless of activation. While M1 polarization was associated with increased CD11c expression (3b), analysis of EV+ cells revealed a predominance of CD11c low macrophages among the EV-positive population (3c), indicating that EV uptake preferentially occurs in CD11c low rather than CD11c high subsets.

As expected, CD206 expression increased upon M2 polarization (3d); however, EV uptake was still substantial in CD206^low cells, particularly in M0 and M1 macrophages, highlighting that full M2 differentiation is not required for efficient EV internalization.

A significantly greater percentage of M1 macrophages were mNG+ (29.4%) compared to M0 and M2 counterparts, corresponding to a ∼1.6-fold increase (Figure 3g). However, the MFI of EV+ cells was similar across all groups (Figure 3f), indicating that while M1 macrophages more frequently associate with EVs, the per-cell uptake amount remains comparable.

**Figure 3.**
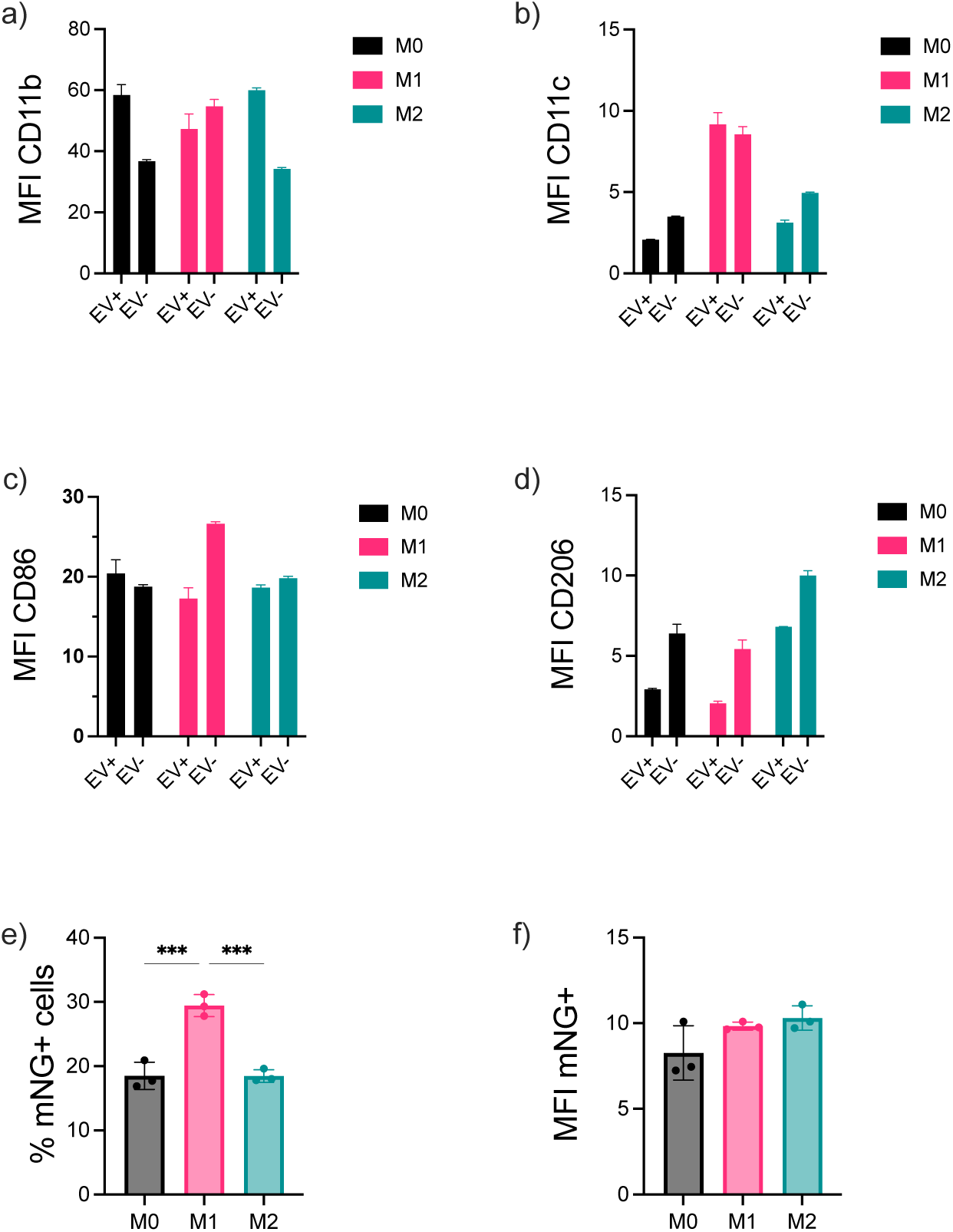
EV uptake by differentiated primary M0 macrophages and polarized M1 and M2 macrophages. **(a)** MFI CD11b expression on EV+ macrophages and untreated macrophages, **(b)** MFI CD11c on EV+ macrophages and untreated macrophages, **(c)** MFI CD86 on EV+ macrophages and untreated macrophages, **(d)** MFI CD206 on EV+ macrophages, and untreated macrophages **(e)** MFI mNG+ on viable cells, **(f)** EV uptake by viable cells as % of mNG+ cells. Macrophages were polarized to M1 and M2 phenotypes according to the Methods section. 1×10^10^ EVs were added for 4 hours. mNG+ cells are EV-positive cells. Cells were stained with F4-80-BV510 – for macrophages, CD11c-Pe/Cy7 for dendritic cells/macrophages, CD11b-APC/Cy7 – myeloid cells, CD4-6-APC – M1 macrophages, and CD206-PE for M2 macrophages. Mean fluorescence intensity (MFI). Black – M0 differentiated macrophages; pink – M1 polarized macrophages, and cyan – M2 polarized macrophages. Statistical differences between multiple groups were determined by one-way ANOVA followed by post-hoc Tukey test. The results represent mean±SD. * represents p<0.05; **, p<0.01; ***, p<0.001 ****, p<0.0001.

Altogether, these findings demonstrate that EV uptake is favoured in CD11b high and CD206 low macrophage subsets, with enhanced uptake frequency observed in M1-polarized cells. This suggests that both macrophage activation state and surface marker profile shape EV interaction dynamics.

### Altered EV association with the immune and non-immune cells under inflammatory conditions

While bioluminescence imaging using NanoLuc-tagged EVs provides valuable insight into global EV biodistribution, it lacks resolution regarding the specific cell types responsible for EV uptake in different physiological contexts. To overcome this limitation, we employed high-parameter flow cytometry to quantify the total abundance of immune and non-immune cells across multiple tissues, and concurrently determined the proportion of CD63-GFP–positive (EV-associated) cells within each population (Figure 4). EVs were administered IV — a clinically relevant route — following acute inflammation induced by LPS.

**Figure 4.**
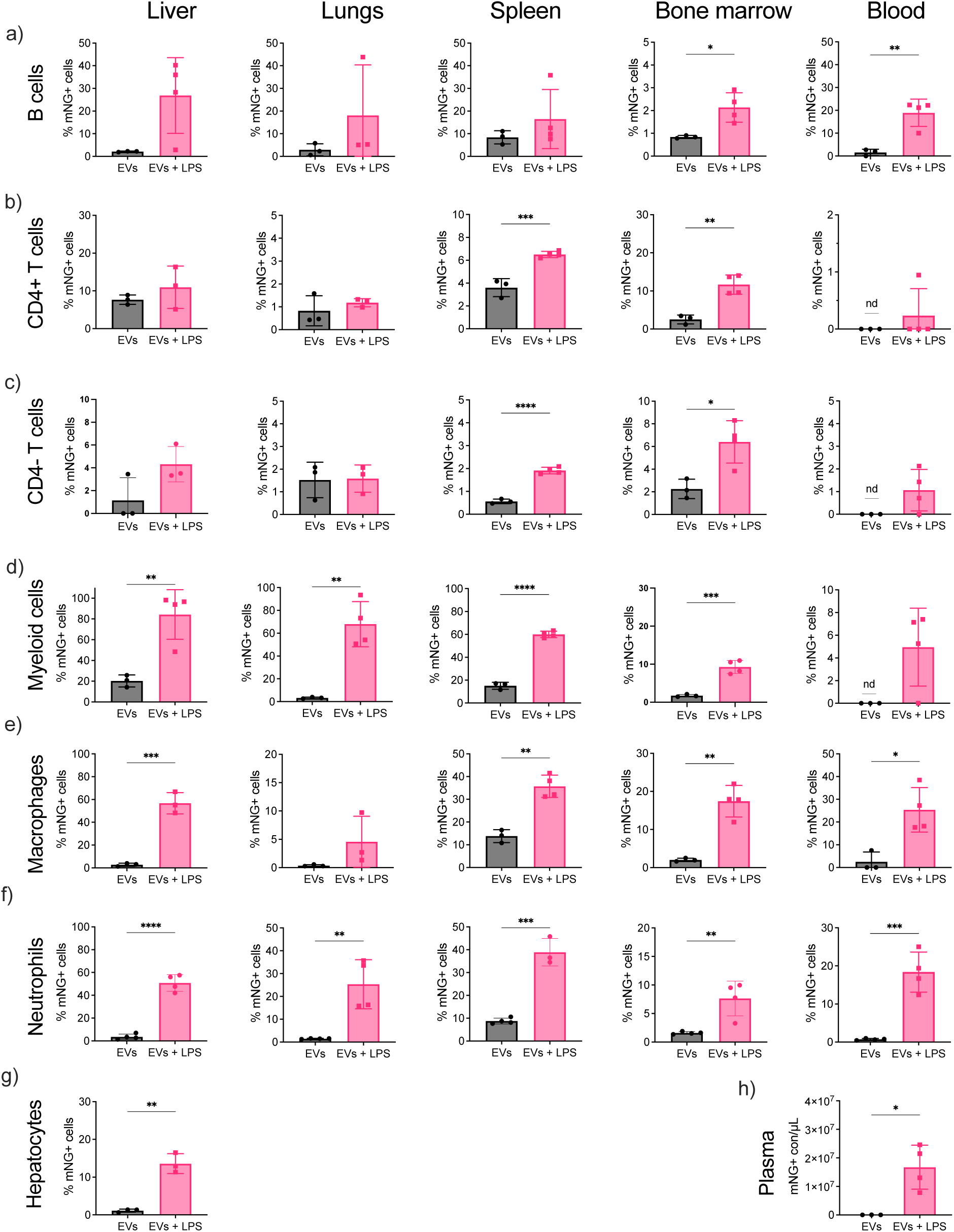
EV association with immune and non-immune cells and biodistribution to various organs. % of EV positive **(a)** B cells; **(b)** CD4 T cells; **(c)** CD4-T cells; **(d)** myeloid cells; **(e)** macrophages; **(f**) neutrophils in the respective tissues: the liver, lungs, spleen, BM, and blood; **(g)** hepatocytes, and **(h)** plasma. Inflammation in mice was induced by a single dose of LPS at the concentration 5 mg/kg I.P. for 4 hours before the administration of EVs. EVs were injected I.V. at 2×10^11^ EVs/dose (n = 3 mice) for 2 hours. A control group (PBS) was used for the normalization of the data (n = 1-2). The immune cell panel was changed depending on the cell type analysis. Cells from each organ were stained for the hematopoietic differentiation marker CD45 (CD45+), macrophages (F4/80+), myeloid (GR1+ /Cd11b+), B cells (B220), CD4 T cells (CD4+/CD3+), CD4-T cells (CD4-/ CD3+), neutrophils (CD11b+/Ly6G+) and evaluated by Flow cytometry. Hepatocytes were gated after excluding immune cells. Cells associated with EVs were mNG positive. The values were normalized to untreated control mice and calculated as % mNG+ cells of viable or mNG+ cell counts/µL of plasma. Black – non-primed healthy mice, pink – LPS-primed mice. ND – non-detectable. Statistical analysis was performed by a two-tailed t-test. * Represents p<0.05, **represents p<0.01, ***p<0.0001 and ****p<0.0001. The results represent mean±SD.

B cells (B220+) (Figure S4, S5) displayed low basal uptake of EVs across tissues in EV injected healthy mice, with the spleen showing the highest association at 8.5%. Under inflammatory conditions, we observed marked increases in EV-positive B cells, most notably in the liver (27%), lungs (18%), and spleen (17%) (Figure 4a). Interestingly, total B cell numbers in these organs remained relatively stable (Figure S6a), suggesting that LPS enhances EV uptake without necessarily expanding the B cell compartment.

CD4+ T cells (Figure S4, S5) showed low to moderate EV uptake under steady-state conditions, with the spleen and BM as primary sites of association. Upon LPS challenge, we observed a 1.8-fold increase in EV-positive CD4+ T cells in the spleen and a 4.6-fold increase in the BM (Figure 4b). Despite this increase in EV association, total CD4+ T cell numbers remained relatively consistent, except for a modest expansion in the spleen and BM under inflammation (Figure S6b).

CD4-T cells (Figure S4, S5), demonstrated tissue-specific increases in EV uptake under inflammatory conditions, most notably a 3.4-fold increase in the spleen and a 2.8-fold increase in the BM (Figure 4c). Notably, EV association remained low in the lungs and blood regardless of LPS priming (Figure S6c), consistent with reduced total CD4-T cell abundance in these tissues.

Myeloid cells (Gr1+CD11b+) (Figure S4, S5) exhibited the most dramatic increase in EV association following LPS administration, despite an overall reduction in their total numbers in key tissues such as the liver, lung, and spleen (Figure S6d). In the liver, the proportion of EV-positive myeloid cells rose sharply from 20.2% in healthy mice to 84.3% in LPS-treated mice—a more than 4-fold increase. Similarly, the lungs showed a 20-fold rise in EV-positive myeloid cells, the spleen a 4-fold increase, and the bone marrow a 5.4-fold increase. Notably, in healthy mice, no EV-positive myeloid cells were detected in the blood, whereas 5% became EV-positive following LPS priming(Figure 4d). These findings suggest that while systemic inflammation enhances EV uptake by myeloid cells, it also leads to a depletion of their overall tissue abundance, possibly due to redistribution, activation-induced cell death, or egress to circulation and inflamed sites.

Macrophages (F4-80+) (Figure S4, S5) (demonstrated selective and robust EV association under inflammation. Only 2.8% of liver macrophages were EV-positive in control mice, compared to 56.7% post-LPS. A similar trend was observed in the lungs, where EV uptake increased 12.6-fold (0.36% to 4.5%), in the BM (8.6-fold), and in the blood (25.4% vs. 2.5%) (Figure 4e). These changes occurred alongside moderate decrease in total macrophage abundance, especially in peripheral tissues (Figure S6e).

Neutrophils (Figure S4, S5) displayed a dramatic increase in EV association in LPS-primed mice, with a 14.2-fold increase in the liver, 17-fold in the lungs, 4.4-fold in the spleen, 4.8-fold in the bone marrow, and a striking 25.9-fold increase in peripheral blood (Figure 4f). However, this enhanced EV uptake was accompanied by a notable reduction in total neutrophil numbers in these tissues (Figure S6f), suggesting that a smaller neutrophil pool becomes more enriched in EV-positive cells under inflammatory conditions. This paradox may reflect neutrophil activation, apoptosis, or trafficking out of tissues in response to LPS and EV exposure.

Non-immune cells (CD45–) (Figure S4, S5), including hepatocytes, also demonstrated enhanced EV uptake in response to LPS. In healthy mice, only ∼1% of hepatocytes were GFP-positive, whereas in LPS-primed animals this rose to 13.6% (Figure 4g). These findings were not explained by changes in hepatocyte number (Figure S6g), suggesting inflammation increases EV association or internalization at the individual cell level.

Finally, to validate these cellular findings, we measured circulating CD63-mNG+ EVs in plasma using single-vesicle imaging flow cytometry. LPS-primed mice showed significantly elevated EV levels compared to controls (Figure 4h), corroborating our previous NanoLuc imaging results and further confirming that systemic inflammation alters both EV biodistribution and cellular engagement.

### EV-Associated Neutrophils Exhibit Elevated CD11b Expression Across Tissues

Given that EV administration induced neutrophil migration, we next investigated whether exogenous EVs also triggered neutrophil activation in target tissues. Neutrophils were identified as CD45⁺Ly6G⁺ cells, and their activation status was assessed by measuring CD11b expression—a complement receptor involved in phagocytosis and a well-established marker of neutrophil activation^32,33^. Since CD11b can facilitate EV uptake, its upregulation may reflect a functional response to EV interaction.

We compared the MFI of CD11b in EV+ and EV− neutrophils from both healthy and LPS-primed mice. In healthy mice, EV+ neutrophils exhibited significantly higher CD11b expression in the lungs, bone marrow, and spleen compared to their EV− counterparts (Figure S7). A similar trend was observed in LPS-treated mice, where CD11b was also significantly upregulated in EV+ neutrophils across these tissues (Figure S7). Notably, in the liver, CD11b upregulation in EV+ neutrophils were only observed in LPS-treated mice, while no significant activation was detected in the liver of healthy controls (Figure S7).

These findings suggest that EV interaction enhances neutrophil activation, with tissue-specific modulation depending on the inflammatory context.

### Enhanced EV uptake lacks correlation with functional delivery of EV protein cargo

A major barrier limiting the therapeutic application of EVs for intracellular protein delivery is their inefficient endosomal escape after uptake. Previous work by others and by our group has consistently demonstrated that although EVs can be efficiently internalized by target cells, the majority of their cargo remains trapped within endosomal compartments, severely restricting functional delivery both *in vitro* and *in vivo*^34–36^

Given that inflammation markedly enhanced EV association with various immune and non-immune cells across tissues, we next investigated whether this increased cellular interaction could improve *in vivo* biodistribution and ultimately translate into more efficient functional delivery. To explore this, we utilized our previously optimized Cre delivery platform, in which Cre recombinase is tethered to CD63 via a pH-sensitive linker^37^ that enables cargo release within endosomal compartments. Functional delivery was assessed using Ai9 Cre-reporter mice, where successful cytosolic delivery and nuclear activity of Cre results in irreversible TdTomato expression.

Despite observing significantly greater EV accumulation and cellular uptake in LPS-treated mice, no meaningful increase in recombination events was detected when compared to untreated controls across major organs, including the liver, lung, spleen, and kidney (Figure S8). These findings suggest that enhanced uptake alone is insufficient to overcome the fundamental barrier of endosomal escape *in vivo*.^37,38^

Additionally, we noted variable background tdTomato fluorescence in both LPS-treated and PBS-injected control groups, particularly in the spleen. This background signal may be attributed to several factors. Systemic inflammation induced by LPS can activate endogenous promoters or stress-responsive pathways, potentially leading to low-level, Cre-independent TdTomato expression. Moreover, the spleen’s high cellular turnover and immune activity might contribute to nonspecific activation of the reporter gene. These observations underscore the importance of including appropriate controls and interpreting reporter gene expression with caution, especially under inflammatory conditions.

### *In Vivo* Mapping of EV-Associated Surface Proteins Highlights Corona Remodeling in Inflammation

The protein corona, known to shape the biological identity and biodistribution of synthetic nanoparticles^39^, similarly forms around EVs upon systemic exposure^14^. We hypothesized that systemic inflammation alters the EV corona, influencing their in vivo fate. Elevated inflammatory proteins such as complement factors and opsonins may bind to EV surfaces and redirect their tissue targeting, providing a mechanistic basis for the altered biodistribution observed under inflammatory conditions^40^.

To investigate how systemic inflammation reshapes the protein corona^41^ of EVs, we employed our previously optimized APEX2-mediated proximity biotinylation strategy to selectively label surface-associated proteins ex vivo on EVs incubated in plasma collected from either untreated mice or mice subjected to LPS-induced systemic inflammation. (Figure 5a)^42^. Across all 12 biological replicates, we identified a total of 3,986 proteins, of which 2,850 high-confidence biotinylated proteins were retained after filtering out cRAP contaminants and red blood cell (RBC)-derived components (Figure S9, S10). Technical reproducibility was high, with each sample yielding approximately 3,900–4,000 proteins, and over 3,000 proteins consistently identified across all samples, confirming robust labelling and mass spectrometry performance (Figure S10a, b, c).

**Figure 5.**
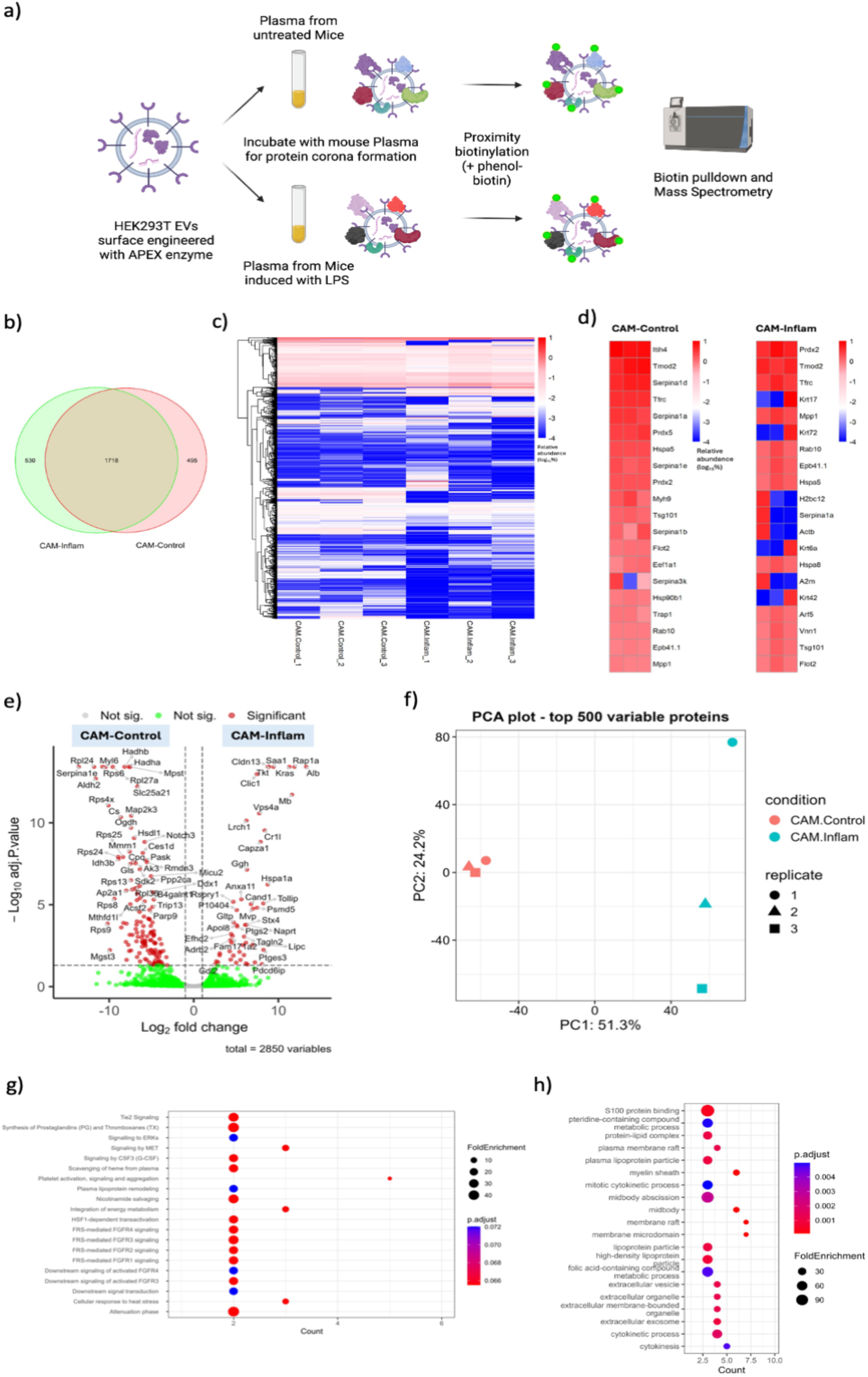
APEX2-based proximity labelling enables characterization of inflammation-specific EV protein corona. **(a)** Schematic of APEX2-based biotinylation workflow. EVs engineered to express APEX2 were incubated with plasma from untreated or LPS-treated mice. Biotinylated proteins were pulled down and analysed by mass spectrometry. **(b)** Venn diagram showing overlap of biotinylated proteins in CAM-Control vs. CAM-Inflam groups. **(c)** Heatmap depicting relative abundance of proteins (log10 scale) after removal of RBC-associated and contaminant proteins. **(d)** Selected proteins enriched in each group, visualized by relative abundance (log10%). **(e)** Volcano plot illustrating differentially enriched proteins (adjusted p < 0.05, log2FC ≥1 or ≤-1) between CAM-Control and CAM-Inflam groups. **(f)** PCA plot of the top 500 variable proteins showing clear separation between inflamed and control EV coronas. (**g-h)** GO and Reactome pathway enrichment analysis of upregulated proteins in CAM-Inflam samples reveals involvement in protein folding, immune regulation, and vesicle-mediated transport processes.

To rule out technical artifacts from erythrocyte contamination, we examined the abundance of canonical RBC proteins. Although trace levels of haemoglobin subunits (Hba, Hbb), Hpx, and Hp were observed, their overall abundance was low and comparable across groups, consistent with mild haemolysis or inflammation-associated RBC-EV shedding (Figure S10d). Moreover, the presence of these proteins may be partially attributed to intrinsic peroxidase activity in RBCs, which can interfere with APEX2-based labelling.^43^ Albumin analysis further revealed a substantial increase in mouse albumin abundance on EVs from LPS-treated mice, consistent with stronger interactions between circulating EVs and plasma proteins under inflammatory conditions (Figure S10e).

Analysis of biotinylated proteins showed that 1,718 were shared between corona in inflamed and control plasma, while 530 and 495 proteins were uniquely enriched in the CD63 ApeX Modified EVs (CAM)-Inflam and CAM-Control groups, respectively (Figure 5b). Heatmap comparisons of relative abundance (log10%) for top-ranking corona proteins illustrated distinct expression patterns: CAM-Control EVs were enriched for classical EV and intracellular proteins (e.g., Tsg101, Hspa5, Flot2), whereas CAM-Inflam EVs displayed a proteome dominated by acute-phase proteins (e.g., A2m, Alb), cytoskeletal regulators (e.g., Krt17, Actb), and inflammation-associated markers (Figure 5c, d). These profiles reflect a pronounced and reproducible remodelling of the EV corona in response to inflammation. Importantly, while global proteomic profiling of mouse plasma is possible, it remains challenging due to the high dynamic range of protein abundance and dominance of a few highly abundant proteins (e.g., albumin, immunoglobulins), which can mask detection of low-abundance but functionally relevant components.

Differential abundance analysis further confirmed this shift, identifying 51 proteins significantly upregulated in CAM-Inflam EVs and 135 in CAM-Control (adjusted p < 0.05, |log2FC| ≥ 1). Among the proteins enriched during inflammation were Saa1, Rap1a, Cr1l, Alb, and A2m, many of which are known to participate in immune signalling, coagulation, and acute-phase responses (Figure 5e). Principal component analysis of the top 500 most variable proteins showed complete separation of inflamed and control samples along PC1, which explained 51.3% of the variance, reinforcing the robustness of the inflammatory signal on corona composition (Figure 5f). Notably, unsupervised hierarchical clustering of the dataset revealed a clear segregation between groups and strong intra-group consistency, highlighting that inflammation drives global remodelling of the EV surface proteome.

Functional enrichment analysis of upregulated proteins in the inflamed group revealed significant overrepresentation of biological processes involved in complement activation, blood coagulation, platelet degranulation, and immune effector regulation. Pathways such as integrin-mediated adhesion and extracellular matrix organization were also enriched. Notably, proteins such as complement C3, alpha-2-macroglobulin (A2m), fibrinogen subunits, haptoglobin, and clustering are known to influence nanoparticle stability, reduce immune clearance, and prolong circulation—suggesting that systemic inflammation reprograms the EV protein corona to favour plasma retention and immune modulation (Figure 5g, h; S11).

Taken together, these results demonstrate that systemic inflammation induces a distinct and reproducible reorganization of the EV protein corona. The shift toward plasma- and immune-related proteins under inflammatory conditions likely reflects both increased exposure to host factors and active recruitment of stabilizing components. While this altered corona composition supports prolonged circulation and altered biodistribution of EVs, it may simultaneously mask intrinsic EV features important for intracellular delivery— highlighting the complex and context-dependent nature of EV behaviour *in vivo*.

## Discussion

This study provides a comprehensive examination of how systemic inflammation alters the biodistribution, cellular targeting, and retention of EVs, shedding light on their interaction with immune and non-immune cells *in vivo*. While prior studies have shown that EVs primarily accumulate in the liver and spleen under physiological conditions, our findings in an LPS-induced inflammation model reveal that systemic inflammation enhances EV accumulation across multiple organs and prolongs their plasma half-life, indicating a suppression or redirection of normal clearance mechanisms. This prolonged circulation increases the window for EV interaction with immune and parenchymal cells in inflamed tissues.

Similar observations were made in the TNBS-induced colitis model, where EVs exhibited enhanced accumulation in inflamed intestinal tissue, further supporting the notion that inflammation actively reshapes EV biodistribution and tissue targeting^44,45^. These findings suggest a conserved response across different models of inflammation, pointing to an inflammation-driven mechanism that facilitates EV access to activated or damaged tissues.

Taken together, these insights highlight not only the dynamic nature of EV pharmacokinetics in disease contexts but also their therapeutic potential in inflammatory disorders. Prolonged circulation and preferential localization to inflamed tissues make EVs an attractive platform for targeted delivery of anti-inflammatory agents. Moreover, these biodistribution shifts may help explain the mechanism of action of endogenous EVs in resolving inflammation, as they can deliver immunomodulatory signals—such as regulatory RNAs or membrane ligands—to key immune effectors like neutrophils, macrophages, or epithelial cells. Leveraging this natural tropism and modifying EV cargo could offer a promising strategy for treating diseases such as colitis, sepsis, or cytokine storm syndromes.

One of the most striking observations was the increased association of EVs with immune cells, particularly in inflamed tissues. In physiological conditions, EVs showed limited interactions with circulating immune cells, with minimal uptake by T cells and low association with myeloid cells. However, under inflammatory conditions, EVs were significantly more likely to interact with immune cells, including a tenfold increase in association with myeloid cells and higher uptake by T cells. Notably, M1-polarized macrophages demonstrated higher EV uptake *in vitro* than M0 or M2 phenotypes, indicating that macrophage polarization plays a critical role in modulating EV internalization. This polarization-dependent uptake suggests that in pro-inflammatory environments, specific immune cell states could influence the retention and clearance of EVs, potentially impacting the efficacy of EV-based therapies.

Additionally, we observed a marked increase in EV association with neutrophils in the bone marrow and peripheral tissues. The recruitment of neutrophils to inflamed organs following EV administration indicates that EVs may play a role in modulating immune cell migration during inflammation. This interaction was accompanied by upregulation of CD11b on neutrophils, a marker of activation, suggesting that EVs may influence not only immune cell distribution but also activation states, which could be harnessed for therapeutic purposes in targeting inflamed tissues.

Interestingly, our data show that EVs also associate with non-immune cells, such as hepatocytes, in inflamed conditions. Nearly 14% of hepatocytes were found to interact with EVs under LPS-induced inflammation, compared to only 1% in physiological states. This association with non-immune cells may contribute to the observed prolonged EV retention in inflamed organs. Non-immune cells lack efficient mechanisms to process and clear EVs, which could lead to increased accumulation and even exacerbate organ dysfunction in inflammatory diseases. Such findings underscore the need to consider both immune and non-immune cell interactions when designing EV-based therapeutic strategies.

Using APEX2 proximity labeling, we identified inflammation-specific enrichment of plasma proteins involved in immune response and vesicle trafficking, suggesting a reshaped corona that may contribute to enhanced tissue retention and cellular uptake. Although EVs demonstrated broader biodistribution and increased cellular association in inflamed tissues, they did not induce functional Cre-mediated recombination in Ai9 reporter mice. This highlights that while systemic inflammation significantly alters the in vivo behaviour of EVs—enhancing tissue accumulation and engagement with specific cell populations— these changes alone are insufficient for effective intracellular delivery, likely due to limited endosomal escape. Importantly, inflammation may prolong EV circulation time and redirect their cellular tropism, favouring uptake by activated immune or endothelial cells. These inflammation-driven shifts in pharmacokinetics and targeting could contribute to the mode of therapeutic action of EVs in vivo, independent of direct cytosolic delivery. Overall, these findings emphasize the need to account for inflammatory status when designing and evaluating EV-based therapies.

## Conclusion

Our findings highlight the profound impact of systemic inflammation on the biodistribution, cellular targeting, and retention of EVs *in vivo*. Inflammatory conditions not only increase the association of EVs with various immune cell types, including B cells, CD4 and CD4-T cells, and neutrophils, but also promote EV interaction with non-immune cells, such as hepatocytes. These enhanced interactions underscore the potential for EVs to modulate immune responses and interact with a broad range of cell types under pathological conditions. However, the limited improvement in intracellular delivery emphasizes the need for further engineering of EVs to enhance endosomal escape and cargo release. These insights into EV dynamics in inflamed environments are essential for the development of targeted EV-based therapies, particularly in chronic inflammatory diseases, where selective cell targeting and modulation of immune responses are critical therapeutic goals.

## Author Contributions

D.G and S.E.A conceived the study. S.P performed majority of the in vivo experiments. D.R.M, X.L, Z.N, S.R and G.Z assisted with the in vivo experiments. A.G and A.M.Z performed the flow cytometry experiments. W.Z, D.Y and K.I performed and analysed the proximity labelling of EVs. E.V.W, Z.N and R.E.V performed the Cre recombinase in vivo experiment in Ai9 mice model. O.P.W, M.P.W and S.E.A provided the resources. S.P and D.G wrote the manuscript with the help from other co authors.

## Acknowledgment

D.R.M and O.P.W are supported by the Swedish Research Council (VR, 2022-02449), the Swedish Cancer Society (Cancerfonden, project 23 2935 Pj), Radiumhemmet (project #241392), the Center for Innovative Medicine (CIMED) junior investigator grant (FoUI-976434), and Karolinska Institutet (2-116/2023). A.Z is supported by the Cancerfonden, D.Y is supported by the Clarendon Fund, Oxford. D.G is supported by BHF Cureheart and MRC Core Therapeutic Genomics. SEA is supported by the European Research Council (ERC) under the European Union’s Horizon 2020 research and innovation program (DELIVER, grant agreement No. 101001374; EXPERT, grant agreement No. 825828), Swedish Foundation of Strategic Research (FormulaEx, SM19-0007), Cancer Foundation (grant No. 21-1762-Pj-01-H), Swedish Research Council (grant No. 4–258/2021) and Brain foundation (Hjärnfonden) contract (FO2024-0073-TK-113)

