## Supplementary Figures for "Systemic Inflammation Modulates Clearance and drives Extra-Hepatic Distribution of Extracellular Vesicles"

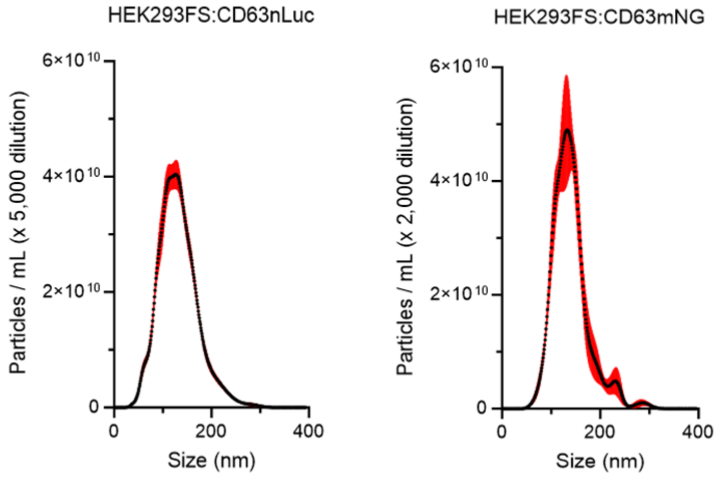


| | |
|---|---|
| Mean | 132 +/- 1.5 nm |
| Mode | 118 +/- 4.4 nm |
| D10 | 85 +/- 0.8 nm |
| D50 | 128 +/- 1.8 nm |
| D90 | 182 +/- 3.1 nm |

| | |
|---|---|
| Mean | 139.4 +/- 1.4 nm |
| Mode | 130.7 +/- 7 nm |
| D10 | 98.3 +/- 2.0 nm |
| D50 | 134 +/- 2.0 nm |
| D90 | 181 +/- 5.2 nm |

a)

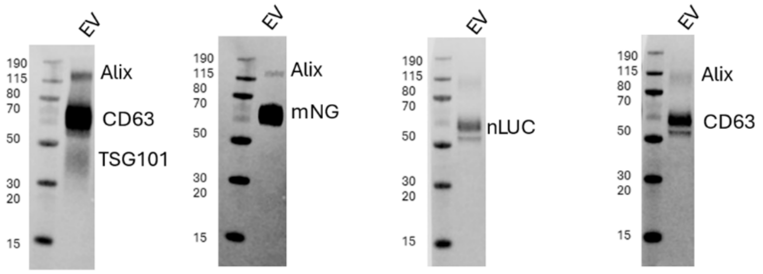


b)

**Figure S1**. **EV characterization by NTA and Western blot. (a)** Particle size distribution and concentration of HEK293FS:CD63NanoLuc and HEK293FS:CD63mNG EVs determined by NTA. **(b)** The WB for CD63 (bound to -mNG or -NanoLuc), mNG (bound to CD63), NanoLuc (bound to CD63), Alix, and TSG101.

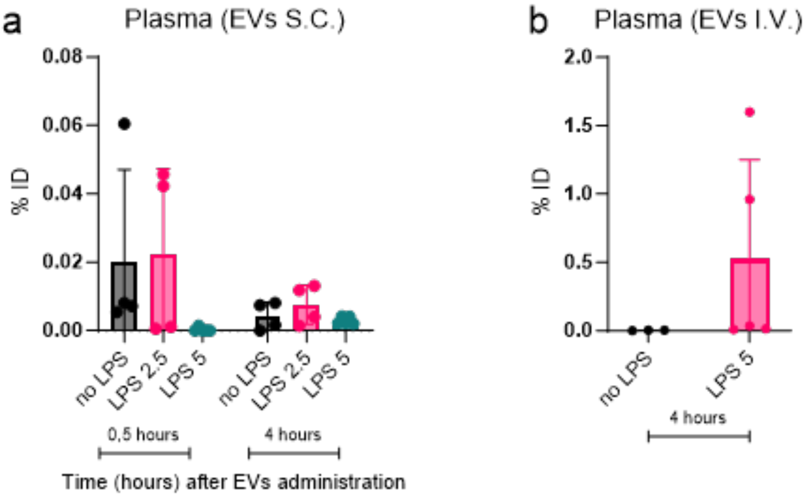


**Figure S2. EV accumulation in plasma is dependent on the administration path and the severity of induced inflammation.** EVs were quantified by the bioluminescent assay as % ID/mL in plasma after **(a)** S.C administration and **(b)** I.V. Inflammation in mice was induced by a single dose of LPS I.P. at the concentration 2.5 mg/kg (LPS 2.5) or 5 mg/kg (LPS 5) for 4 hours before the administration of EVs. EVs were injected at $1X10^{11}$ EVs/dose (n = 3 – 5 mice). Blood was collected at 30 minutes (only S.C.) and 4 hours of post-EV administration and plasma was prepared according to the Methods section. The control group (n = 2) was used to normalize the data. Black – non-primed healthy mice, pink – LPS-primed mice with LPS 2.5 mg/mL, cyan – LPS-primed mice with LPS at 5 mg/kg. Statistical analysis was performed by one-way ANOVA. The results represent mean ± SD.

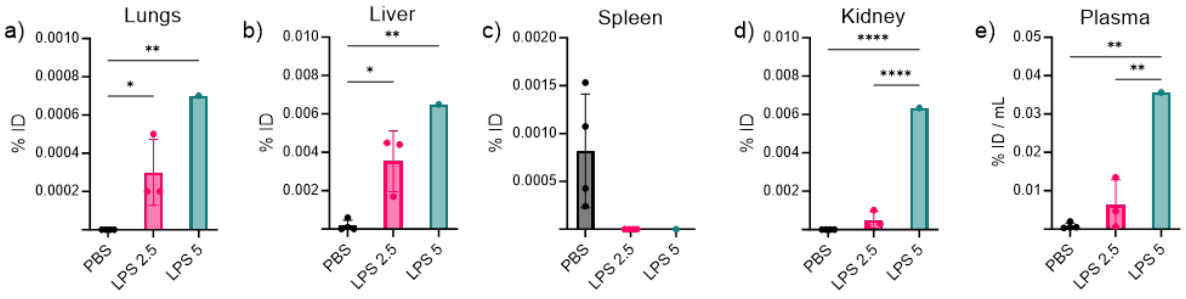


**Figure S3. EV biodistribution depended on the administration path and the severity of induced inflammation.** EVs were quantified by the bioluminescent assay as % ID (organ/mouse) in **(a)** lungs, **(b)** liver, **(c)** spleen, **(d)** kidney, and as % ID/mL in **(e)** plasma.

Inflammation in mice was induced by a single dose of LPS at the concentration of 2.5 mg/kg (LPS 2.5) or 5 mg/kg (LPS 5) I.P. for 4 hours before the administration of EVs. EVs were injected S.C. at 1X10$^{11}$ EVs/dose (n = 3 – 5 mice). Tissues were collected 24 hours post-EV administration. The control group (n = 2) was used for the normalization of the data. Black – non-primed healthy mice, pink – LPS-primed mice with LPS 2.5, cyan – LPS-primed mice with LPS 5. Statistical analysis was performed by one-way ANOVA. * represents p<0.05; **, p<0.01; and ****, p<0.0001. The results represent mean ± SD.

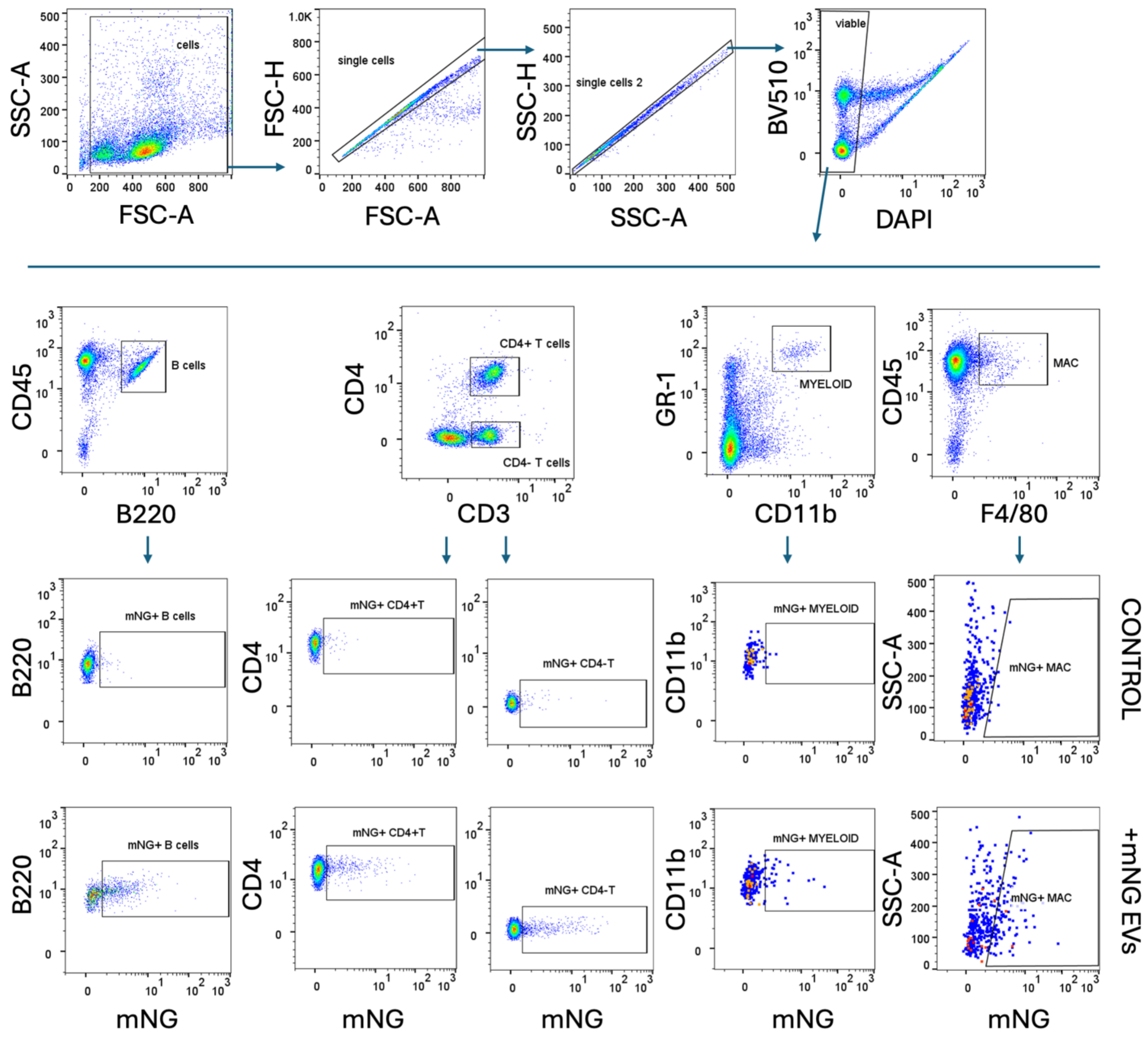


**Figure S4. Representative example of flow cytometry gating scheme to identify immune cells and EV uptake by immune cells** (B cells, CD4 T cells, CD4- T cells, myeloid cells, and macrophages) in the spleen.

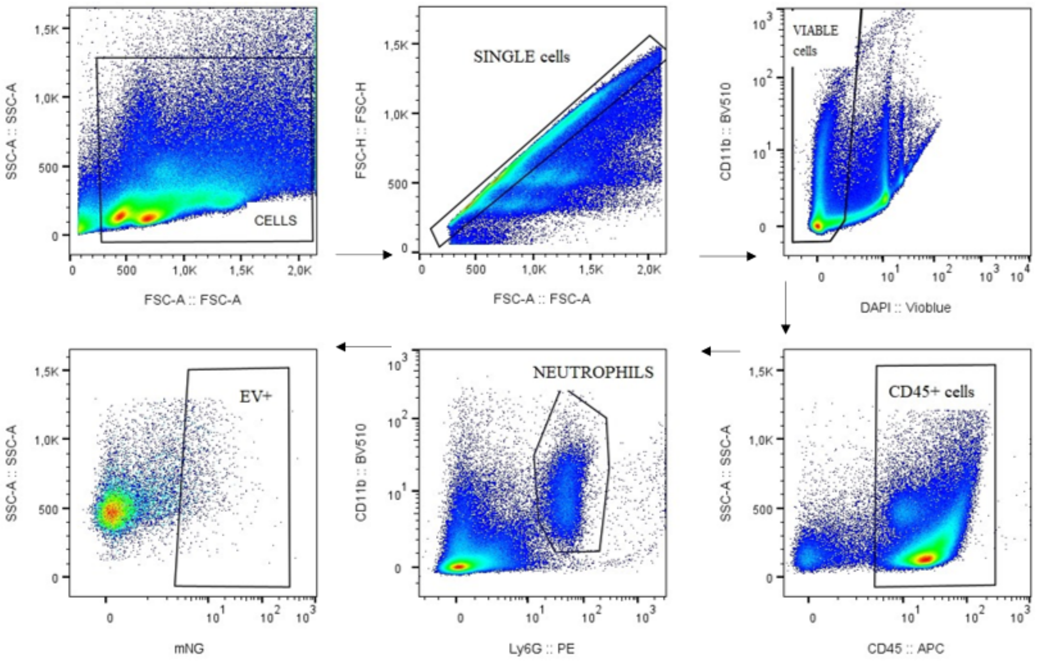


**Figure S5. Representative example of flow cytometry gating for neutrophils in the spleen.**

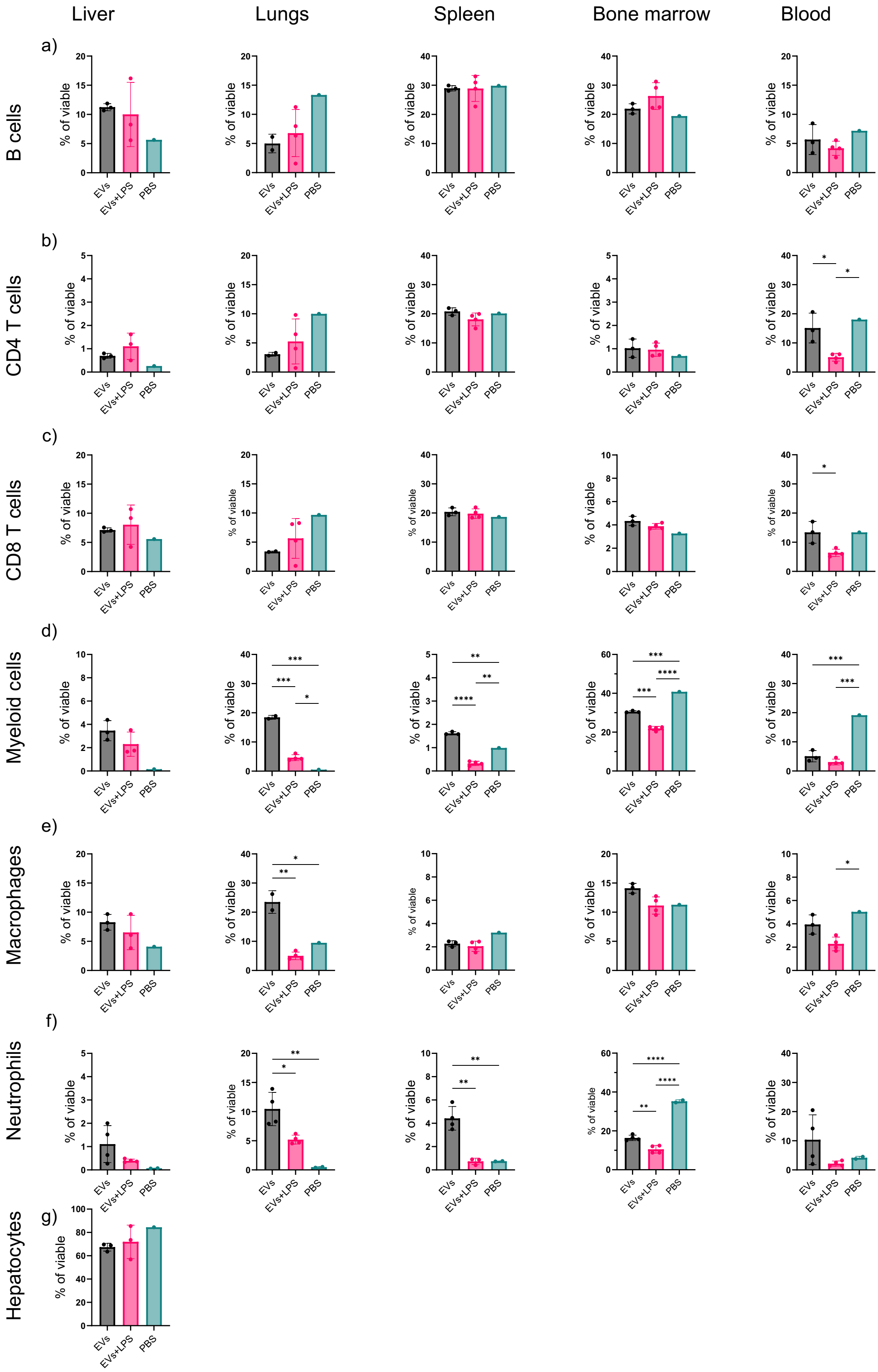


**Figure S6. Immune cells proportion in various tissues**. Supplementary figure complementing Figure 4. The proportion of immune cells: **(a)** B cells, **(b)** CD4 T cells, **(c)** CD4-

T, **(d)** myeloid cells, **(e)** macrophages, **(f)** neutrophils as % of viable in the liver, lungs, liver, BM, and blood. g) hepatocytes. Cells were stained for a general immune marker (CD45), macrophages (F4/80), myeloid (GR1/CD11b), B cells (B220), CD4 T cells (high CD4/CD3), CD4- T cells (medium CD4/ CD3), neutrophils (CD11b, Ly6G-PE). Hepatocytes were gated after excluding immune cells. Statistical analysis was performed one-way ANOVA. * represents p<0.05, **represents p<0.01, ***p<0.0001 and ****p<0.0001. The results represent mean±SD.

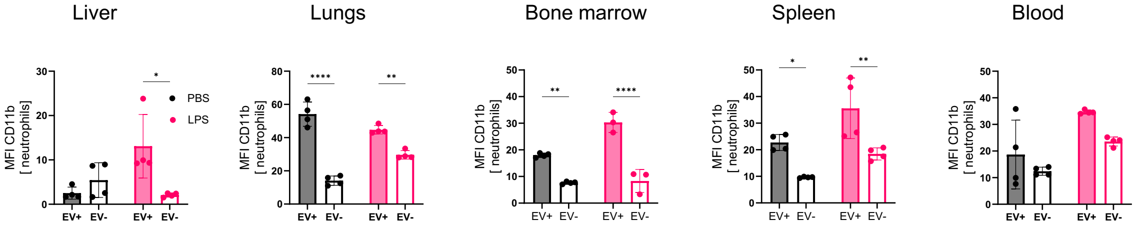


**Figure S7. Upregulation of CD11b+ neutrophils in control and LPS-primed mice in the liver, lungs, spleen, bone marrow, and blood.** Data for analysis were used from the same test as in Figure 4. Neutrophils were gated on CD45-APC, and Ly6G-PE. The expression level of CD11b in neutrophils was determined as MFI CD11b on EV+ neutrophils or EV– neutrophils. EV+ cells are mNG+ cells. Black – non-primed healthy mice, pink – LPS-primed mice; filled columns represent EV+ cells, empty columns represent EV- cells. Statistical analysis was performed by a Two-way ANOVA. * Represents p<0.05, **represents p<0.01, and ****p<0.0001. The results represent mean±SD.

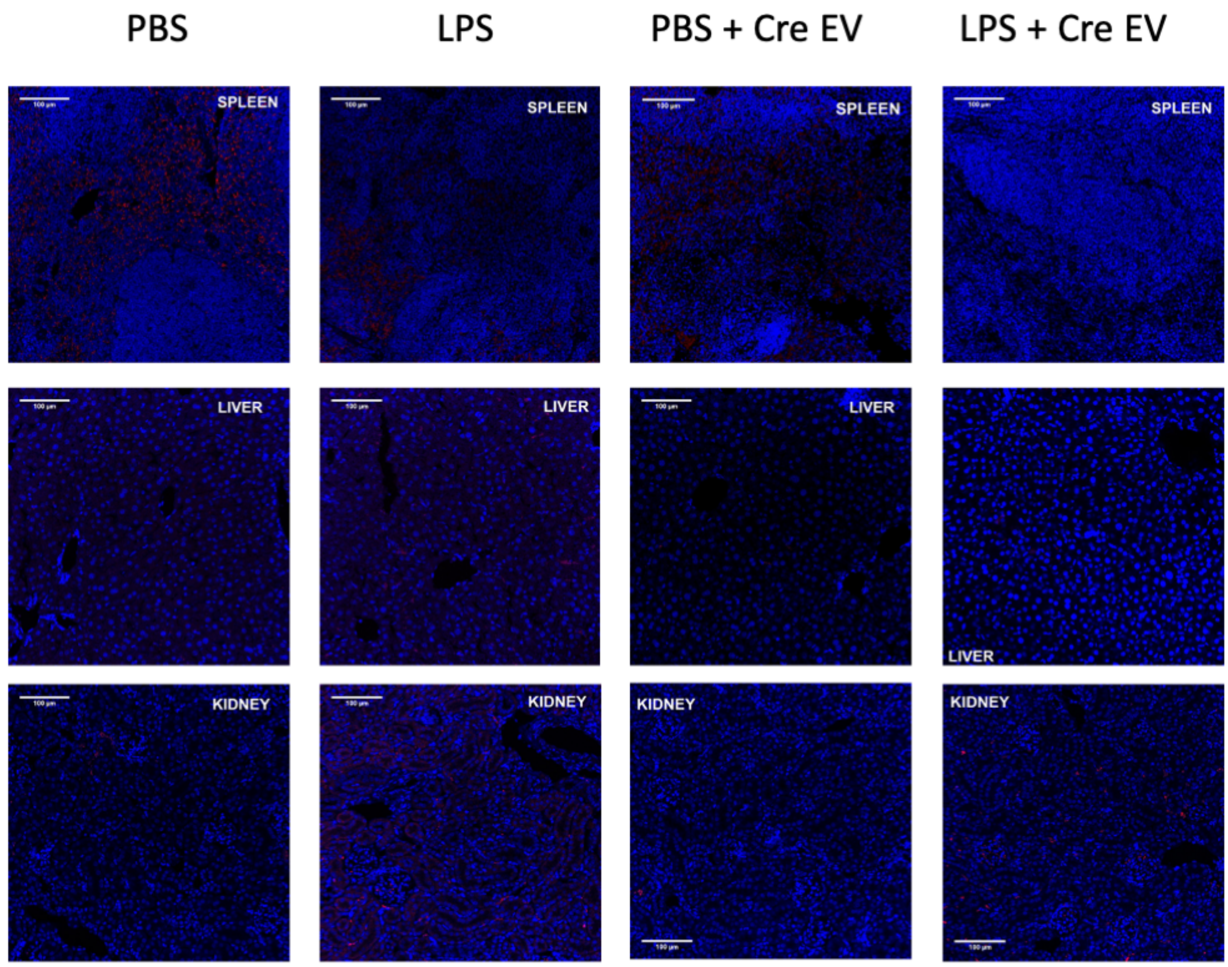


**Figure S8. *In vivo* evaluation of EV-mediated functional protein delivery using Ai9 Cre-reporter mice.** Representative fluorescence images of spleen, liver, and kidney sections collected 7 days post-injection from Ai9 mice under four treatment conditions: PBS only, LPS only, PBS + $1X10^{12}$ CD63 Intein Cre EVs, and LPS + $1X10^{12}$ CD63 Intein Cre EVs. TdTomato fluorescence (red) indicates Cre-mediated recombination; DAPI (blue) marks nuclei. No significant TdTomato expression was detected in any tissue, including in the EV-treated groups, despite increased EV uptake under inflammatory conditions. Notably, some background red fluorescence was observed in spleens of PBS- and LPS-treated controls, consistent with reported tissue autofluorescence or basal reporter leakiness. Scale bar: 100 µm.

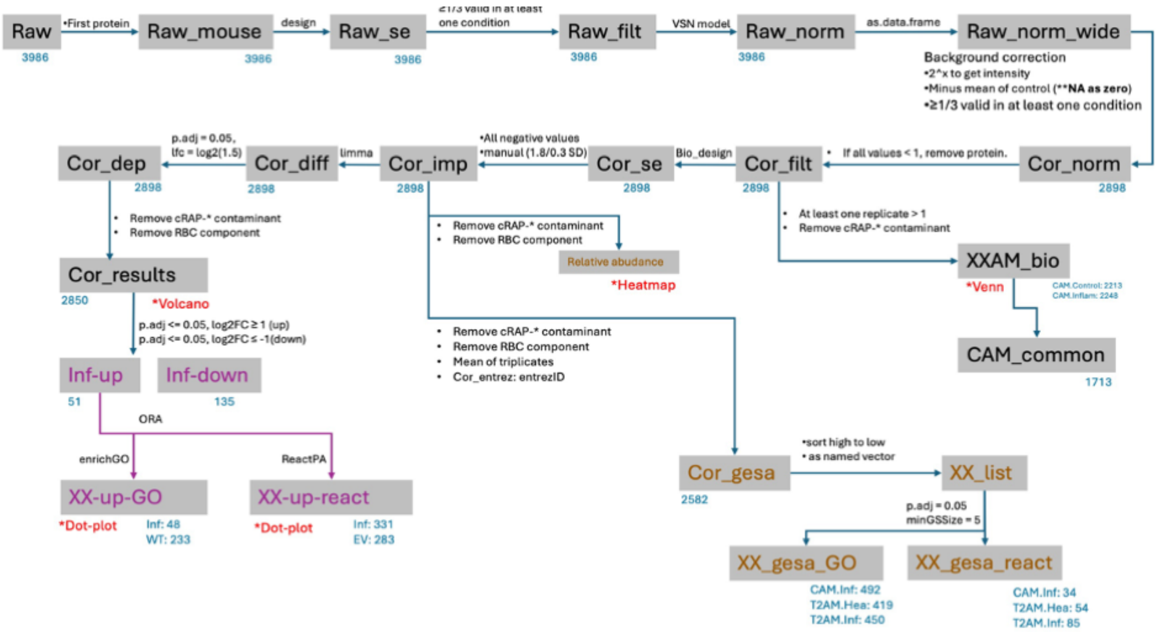


**Figure S9. Workflow for proteomic analysis of EV protein corona using APEX labeling and in vivo plasma interaction.** Schematic overview of the proteomic analysis pipeline. Raw proteomic data were filtered, normalized, and imputed to remove common contaminants and red blood cell components. Multiple downstream analyses, including differential expression (Cor_diff), protein enrichment (ORA and ReactomePA), and gene set enrichment analysis (GSEA), were performed on curated datasets to identify inflammation-specific and EV-specific proteins and pathways.

a)

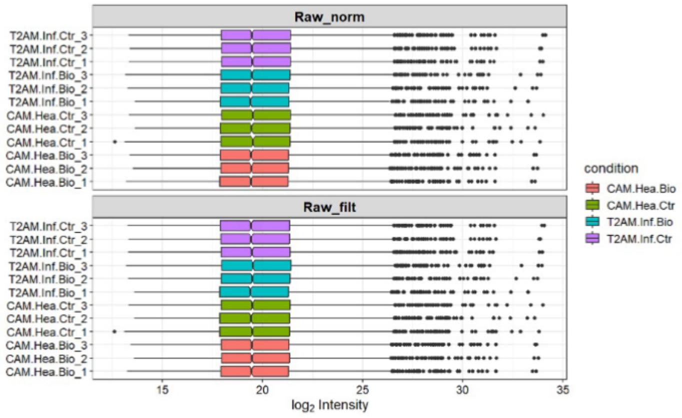

Raw_norm
Raw_filt
T2AM.Inf.Ctr_3
T2AM.Inf.Ctr_2
T2AM.Inf.Ctr_1
T2AM.Inf.Bio_3
T2AM.Inf.Bio_2
T2AM.Inf.Bio_1
CAM.Hea.Ctr_3
CAM.Hea.Ctr_2
CAM.Hea.Ctr_1
CAM.Hea.Bio_3
CAM.Hea.Bio_2
CAM.Hea.Bio_1
15
20
25
30
35
log2 Intensity
condition
CAM.Hea.Bio
CAM.Hea.Ctr
T2AM.Inf.Bio
T2AM.Inf.Ctr


b)

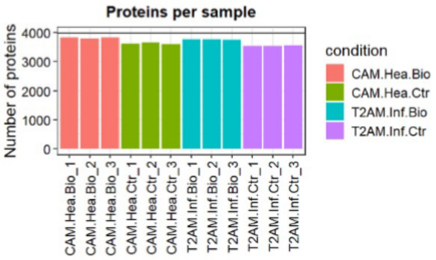

Proteins per sample
Number of proteins
0
1000
2000
3000
4000
condition
CAM.Hea.Bio
CAM.Hea.Ctr
T2AM.Inf.Bio
T2AM.Inf.Ctr
CAM.Hea.Bio_1
CAM.Hea.Bio_2
CAM.Hea.Bio_3
CAM.Hea.Ctr_1
CAM.Hea.Ctr_2
CAM.Hea.Ctr_3
T2AM.Inf.Bio_1
T2AM.Inf.Bio_2
T2AM.Inf.Bio_3
T2AM.Inf.Ctr_1
T2AM.Inf.Ctr_2
T2AM.Inf.Ctr_3


c)

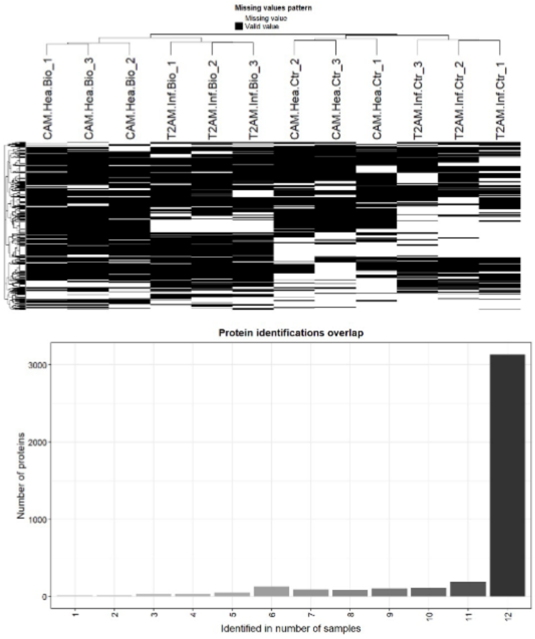

Missing values pattern
Missing value
Valid value
CAM.Hea.Bio_1
CAM.Hea.Bio_3
CAM.Hea.Bio_2
T2AM.Inf.Bio_1
T2AM.Inf.Bio_2
T2AM.Inf.Bio_3
CAM.Hea.Ctr_2
CAM.Hea.Ctr_3
CAM.Hea.Ctr_1
T2AM.Inf.Ctr_3
T2AM.Inf.Ctr_2
T2AM.Inf.Ctr_1
Protein identifications overlap
Number of proteins
0
1000
2000
3000
1
2
3
4
5
6
7
8
9
10
11
12
Identified in number of samples


d)

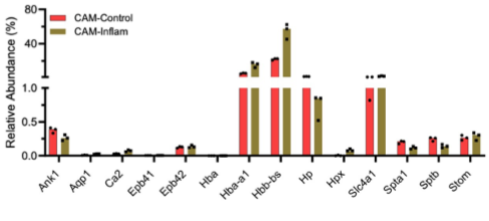

Relative Abundance (%)
CAM-Control
CAM-Inflam
80
40
1.0
0.5
0.0
Ank1
Aqp1
Ca2
Epb41
Epb42
Hba
Hba-a1
Hbb-bs
Hp
Hpx
Slc4a1
Spta1
Sptb
Stom


e)

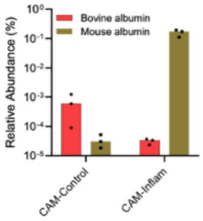

Relative Abundance (%)
Bovine albumin
Mouse albumin
CAM-Control
CAM-Inflam

**Figure S10. Technical validation and data quality control of proteomic dataset. (a)** Boxplots showing intensity distribution across replicates before (Raw_norm) and after (Raw_filt) filtering. **(b)** Bar plot representing the total number of proteins identified per sample across different experimental groups. **(c)** Top: Missing value patterns across samples (black = valid value, white = missing); Bottom: Histogram showing overlap of protein identification across replicates. Most proteins were consistently identified in all samples. **(d)** Relative abundance of red blood cell (RBC)-associated proteins following removal of cRAP contaminants, confirming residual hemoglobin enrichment in CAM-Inflam samples. **(e)** Relative abundance of mouse vs. bovine albumin in CAM-Control and CAM-Inflam samples. Inflammation leads to elevated plasma-derived albumin association with EVs, potentially due to increased vascular permeability and intrinsic peroxidase activity in RBCs enhancing oxidative labeling.

a)

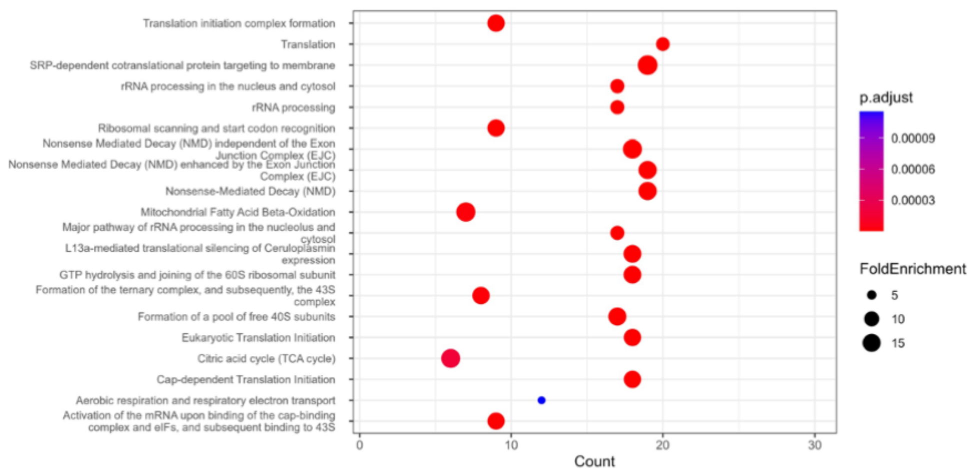


b)

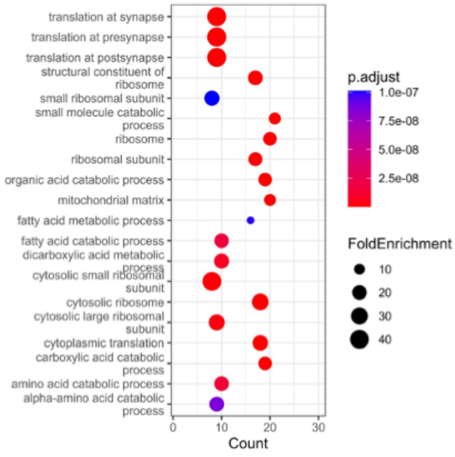


**Figure S11. Functional annotation of inflammation-specific EV corona proteins. (a)** Reactome pathway enrichment analysis of proteins enriched in CAM-Inflam EVs, highlighting pathways related to translation, mitochondrial function, and protein targeting. **(b)** GO biological process analysis showing significant enrichment of terms related to metabolic processes, oxidative stress response, and cytoplasmic translation.
